# Human models of GLE1-associated LCCS1 reveal neural crest deficiency and multisystem developmental failure accompanied by altered RNA metabolism

**DOI:** 10.64898/2026.05.28.726124

**Authors:** Tomáš Zárybnický, Otto J.M Mäkelä, Athina Balatsou, Sule R Yalcin, Sonja Lindfors, Saana Metso, Rocio Sartori Maldonado, Pavlína Gregorová, Juho Väänänen, Laura Rekonen, Kateřina Bůtorová, Peter L Sarin, Kirmo Wartiovaara, Henna Tyynismaa, Niklas Pakkasjärvi, Satu Kuure

## Abstract

Lethal congenital contracture syndrome 1 (LCCS1) is a neurodevelopmental disorder caused by *GLE1* c.432-10A>G variant and presenting fetal akinesia, defects in anterior horn spinal cord, skin, skull, and skeletal muscle development. The uniform prenatal lethality of LCCS1 limits access to patient material, thereby hindering mechanistic studies in physiologically relevant models. To overcome this, human embryonic stem cells (hESCs) carrying the LCCS1 variant, patient-derived fetal fibroblasts, and transcriptomic and proteomic profiling were utilized to examine early GLE1 dysfunction in human cells and tissues. Across LCCS1 cell types, reduced global transcription and translation were observed, while nucleocytoplasmic poly(A)+ RNA distribution was unchanged. Despite its context-dependent effects on proliferation, LCCS1 variant altered mRNA decay kinetics and increased stress granule formation in differentiated cells. LCCS1 hESCs retained core pluripotency but reduced choline acetyltransferase and β tubulin III levels, together with increased neurofilament inclusion incidence, indicate functional immaturity in differentiated spinal motor neurons. Differentiation of LCCS1 hESC-derived gastruloids uncovered broad perturbations in neuromuscular and neural crest derivative development, results which were supported by phenotypes detected in ectodermal organoids and fetal LCCS1 tissue. These findings provide new mechanistic insight into LCCS1 pathogenesis and establish a robust human model framework for studying neurodevelopmental disorders.

## INTRODUCTION

Lethal congenital contracture syndrome 1 (LCCS1, OMIM #253310, autosomal recessive) is a uniformly fatal form of arthrogryposis and neurodevelopmental disorder that manifests with the fetal akinesia deformation sequence, severely atrophic skeletal muscles, and a characteristic developmental defect of the spinal cord (1). The condition becomes apparent around the 13^th^ gestational week and results in intrauterine death before the 32^nd^ week of pregnancy (2). Hallmark findings include degeneration of the anterior horn and descending tracts. Abnormalities such as arthrogryposis and pulmonary and skeletal muscle hypoplasia likely result from akinesia sequence, while hydrops, thin and pale skin, micrognathia, and skull deformities remain consistent findings in affected fetuses, although their underlying pathogenesis remains elusive.

LCCS1, lethal arthrogryposis with anterior horn cell disease (LAAHD) and certain familial forms of amyotrophic lawteral sclerosis (ALS) arise from pathogenic variants in human *GLE1* gene (3–5). Although LAAHD presents with a milder phenotype, both LCCS1 and LAAHD represent severe autosomal recessive motor neuron diseases with distinct *GLE1* variants (6), leading to intrauterine (LCCS1) or early neonatal (LAAHD) death. By far, LCCS1, caused by a homozygous A to G substitution (c.432-10A>G), represents the most severe end of *GLE1* disorders (6, 7). This variant generates an illegitimate splice acceptor site in the intron 3 and produces a nine-nucleotide insertion into *GLE1* mRNA, giving rise to three extra amino acids (proline, phenylalanine and glutamine; T144_E145insPFQ) in the human GLE1 protein (6). The insertion of three extra amino acids is predicted to lay in the coiled-coil domain of GLE1 protein, possibly disturbing its homo-oligomerization (6, 8).

Previous studies of GLE1 function and binding partners have established its multifunctional role as an essential regulator of DEAD-box proteins (particularly DDXs in mammalian cells) that participates in regulation of multiple phases of the mRNA life cycle, including mRNA export, translation, and stress granule formation (9–12). The first experiments addressing the functional consequences of the *GLE1* c.432-10A>G variant were performed *in vitro* by overexpressing tagged pathogenic GLE1 protein in HeLa and yeast cells (11). These studies suggested that disturbed mRNA export results in nuclear mRNA accumulation in LCCS1 cells and causes the cellular defects of the LCCS1 pathology. Interestingly, knockdown of *GLE1* in HeLa cells not only impairs RNA transport, but also reduces nascent protein translation and increases the stress granule count in cells experiencing challenging conditions (10).

*In vivo* functional studies of *Gle1* deficiency were initially reported in zebrafish (13, 14) and only recently studied in mouse (15). These established the requirement of intact GLE1 function for motor neuron development and differentiation. Unlike in zebrafish, mice with *Gle1* inactivation die before gastrulation, emphasizing the species-specific requirements for RNA biology. This is further underlined by our findings in the *Gle1* knockin (KI) mouse model harboring the nine extra nucleotides at the intron 3/exon boundary of mouse *Gle1* gene (15). Motor neuron development and axonal arborizations are affected in KI embryos but do not cause functional consequences in adult mice. Instead, we identified additional defects in KI mouse sympathetic heart innervation, likely causing their sudden deaths in adulthood. KI cells demonstrated disturbed cell cycle progression, diminished transcription and translation, and dysregulated micro-RNA expression under stressing conditions.

The early fetal death of LCCS1 patients causes quick tissue necrosis, hindering many molecular analyses and complicating the use of patient material in biomedical studies. To tackle this, we utilized human embryonic stem cell (hESC)-derived organoids, patient fetal fibroblasts, and multiomic profiling to identify physiologically relevant GLE1 functions affected by c.432-10A>G variant. Our findings denote previously unknown multiorgan deficits in LCCS1 and expand the importance of mRNA transport to the neuromuscular and neural crest derivative differentiation.

## RESULTS

### Human embryonic stem cells with homozygous pathogenic GLE1 variant of LCCS1 maintain pluripotency but show diminished self-renewal

To study molecular and cellular changes in a relevant context, we employed hESCs and LCCS1 fetus-derived fibroblasts (**Fig. 1**). We exploited CRISPR/Cas12a genome editing to introduce the homozygous *GLE1*_FinMajor_ (c.432-10A>G) variant into H9 hESC line (referred LCCS1-hESCs) (**Fig. S1A**). To validate that the modified *GLE1* gene undergoes alternative splicing as previously described in LCCS1 patients (6), we performed long-read PacBio SMRT sequencing. This confirmed the exclusive detection of the desired nine nucleotide insertion in LCCS1-hESCs (**Fig. 1A and S1B**). Next, overall proliferation and specifically mitotic events were quantified in wild-type (WT) and LCCS1-hESCs. This demonstrated diminished cellular propagation and disturbed cell cycle progression in LCCS1-hESCs, as discerned by 5-ethynyl 2’-deoxyuridine (EdU) labelling the proliferating cells, and by the increase in phospho-histone H3 (Ser10) (pHH3)-positive mitotic nuclei (**Fig. 1B-C, S1C**).

**Figure 1.**
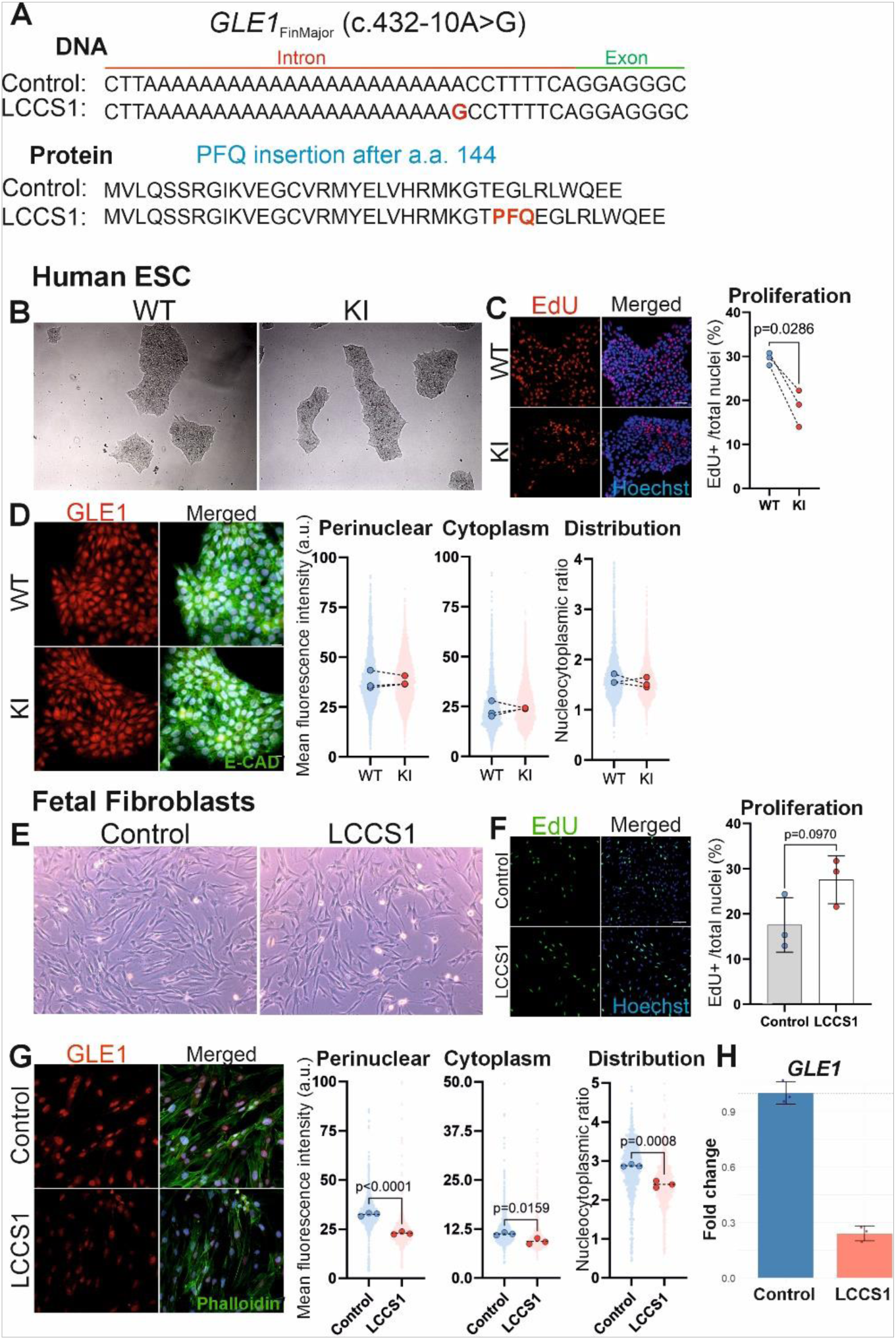
Cell proliferation and GLE1 distribution in human lethal congenital contracture syndrome 1 cells. **A)** Schematics of the *GLE1*_FinMajor_ (c. 432-10A>G) substitution in LCCS1 patient genome (top) and PFQ amino acid insertion as an outcome of *GLE1*_FinMajor_-induced alternative splicing (bottom). **B)** Representative light-microscopy images of wild type (WT) and knockin (KI) H9 human embryonic stem cell (hESC) colonies carrying the *GLE1*_FinMajor_ (c. 432-10A>G) variant. **C)** Cell proliferation assayed in wild type (WT) and knockin (KI) H9 human embryonic stem cell (hESC) by quantification of 5-ethynyl-2′-deoxyuridine (EdU) uptake (10 µM EdU, 2 hours). Representative images shown for wildtype (WT) and LCCS1 knockin (KI) hESCs. Quantification of proliferative cells positive for EdU (red) among all nuclei visualized by Hoechst (blue). The percentage of EdU-positive cells in each independent experiment (over 3000 cells per experiment, three independent experiments) is presented. Scale bar: 50 µm. Paired t-test. **D)** Immunofluorescent analysis of GLE1 (red) protein abundance and distribution in wildtype (WT) and knockin (KI) hESCs, co-stained with E-CADHERIN (E-CAD, green) and Hoechst (blue). Individual data points generated from each cell are plotted (1000 cells per experiment, three independent experiments) and median values from each experiment (large dots) are shown. Scale bar: 20 µm **E)** Representative light-microscopy images of fetal fibroblasts derived from healthy controls and LCCS1 patients. **F)** Cell proliferation assayed in control and patient fibroblasts by quantification of 5-ethynyl-2′-deoxyuridine (EdU) uptake (10 μm EdU, 1 h). Mean ± SD are plotted and percentage of EdU-positive cells in each control and patient (three patients per genotype, over 2500 cells analyzed per patient) presented. Scale bar: 100 μm, two-tailed Student’s t-test. **G)** Immunofluorescent analysis of GLE1 (red) protein abundance and distribution in control and patient fibroblasts, co-stained with phalloidin (green). Individual data points generated from each cell are plotted (400 cells per patient, three patients per genotype), as well as median values from each embryo (large dots). Scale bar: 50 μm. **H)** RNA sequencing analysis of *GLE1* expression in healthy control and LCCS1 patient fibroblasts reveal significantly reduced expression in patient cells.

Since the LCCS1 variant affects cellular proliferation of hESCs, we next assessed whether it also influences their pluripotency. RT-qPCR analysis denoted equal expression of stemness markers *NANOG*, *SOX2*, and *OCT3/4* in cells with and without the variant, while analysis of SOX2 and OCT3/4 proteins revealed equal distribution and abundance (**Fig. S1D-E**). This indicates that the pluripotency of hESCs remains unaffected despite the pathogenic variant causing LCCS1. Interestingly, *GLE1* itself shows significantly lower mRNA expression in the LCCS1-hESCs compared to WT, which does not translate into changes in GLE1 protein abundance or subcellular distribution (**Fig. 1D, S1F**). These results prompted us to study the GLE1 protein stability in LCCS1-hESCs by the thermal shift stability assay. This demonstrated that while GLE1 is generally rather stable, it aggregated at lower temperatures in LCCS1-hESCs (T_agg_ = 78.6 ° C) than in WT (T_agg_ = 84.0 ° C) hECSs (Fig. S1G). Despite maintaining protein levels, this suggests compromised stability of GLE1 in LCCS1-hES cells.

### LCCS1 patient-derived fibroblasts have less GLE1 protein but proliferate more than control cells

Given that the *GLE1* c.432-10A>G variant significantly reduced the cellular proliferation of pluripotent LCCS1-hESCs, we sought to analyze its effects on differentiated patient cells. Microscopy observation followed by EdU labelling demonstrated that, unlike in undifferentiated hESCs, the LCCS1 variant induces increased *in vitro* proliferation of fetus-derived cells compared to control fibroblasts (**Fig. 1E-F**).

Analysis of GLE1 mRNA and protein levels in LCCS1 fetus-derived and control fibroblasts demonstrated that, while all patient cells expressed GLE1 protein, its abundance and mRNA expression were significantly reduced (**Fig. 1G-H**). Immunofluorescence staining for GLE1 demonstrated weaker labeling, particularly at the perinuclear region, leading to a redistribution of the signal and a reduced nucleocytoplasmic ratio. This data suggests different mechanisms for impaired GLE1 functions in stem cells and differentiated fibroblasts: the first are affected by the protein instability and the latter by the diminished protein amount.

### LCCS1-hESC-derived spinal motor neurons display subtle changes indicative of lineage confusion

To investigate the differentiation of LCCS1-hESCs, we first utilized the well-established spinal motor neuron induction protocols, which have successfully been used for modeling different types of motor neuron deficiencies (16, 17). The protocol, spanning over 32 days, yields mature motor neuron-like cells (18). During the stepwise motor neuron differentiation, the first sampling at day 11 (D11) addresses the induction and determination of postmitotic motor neuron progenitor identity (**Fig. 2A**). The induced cells were first characterized by immunostaining with antibodies specific for transcription factors (ISL1/2, OLIG2) necessary for motor neuron lineage induction, progenitor cell differentiation, and regulatory dynamics (19, 20). This showed comparable differentiation of motor neuron progenitors in control and LCCS1-hESCs (**Fig. 2B-C**). To further dissect the *in vitro* neural differentiation, we stained the induced LCCS1-hESCs with HB9 (more committed motor neuron progenitors) and TUJ1 (early neuronal marker) (21). Interestingly, while the TUJ1 signal was comparable between the genotypes, the LCCS1 postmitotic motor neuron progenitors displayed increased HB9 intensity at D11 (**Fig. 3A**). At this point of differentiation, both genotypes presented similar GLE1 signal and distribution (**Fig. S2A**). Together, these results indicate that LCCS1 does not disrupt early motor neuron specification but leads to altered postmitotic motor neuron progenitor identity or maturation, consistent with a defect that emerges after lineage commitment.

**Figure 2.**
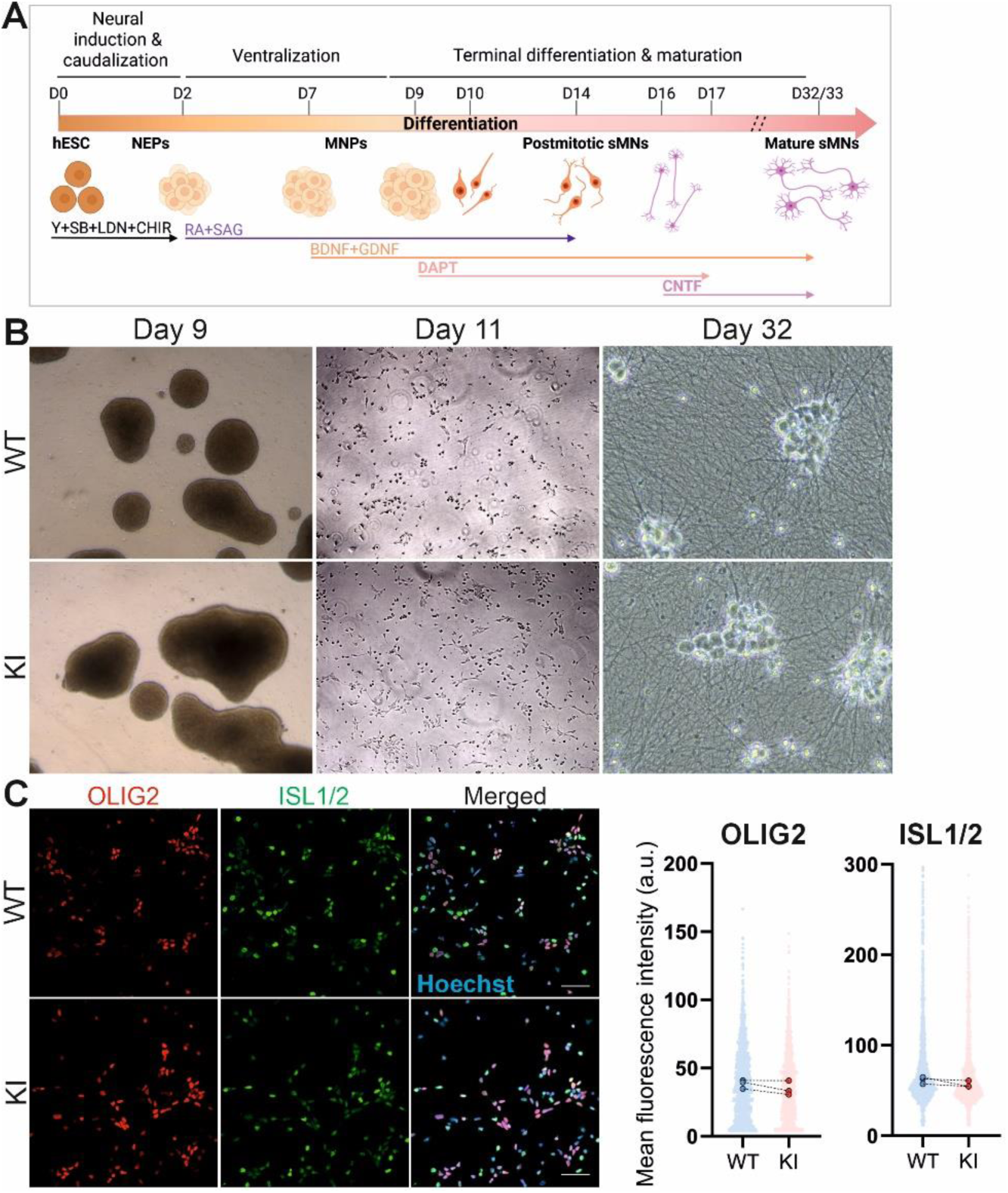
Differentiation of human embryonic stem cells into motor neuron progenitors. **A)** Schematic illustration of motor neuron differentiation protocol. Abbreviations: D, day; hESCs, human embryonic stem cells; NEPs, neuroepithelial stem cells, MNPs, motor neuron progenitors, sMNs spinal motor neurons; Y, Y-27632; SB, SB 431542; LDN, LDN-193189; CHIR, CHIR99021; RA, retinoic acid; SAG, smoothened agonist; DAPT, a γ-secretase inhibitor, BDNF, brain-derived neurotrophic factor; GDNF, glial cell line-derived neurotrophic factor; CNTF, ciliary neurotrophic factor. **B)** Brightfield images of human embryonic stem cells (hESCs) differentiation towards spinal motor neurons (MNs). MN progenitor spheres are shown before dissociation on day nine (D9), illustrations of initiation of postmitotic MNs differentiation (D11) and mature MNs (D32). Scale bar: 200µm **C)** Immunostaining of MN progenitors at D11 of the differentiation with OLIG2 (red) and ISL1/2 (green). Individual data points generated from each cell are plotted (500 cells per differentiation, three independent differentiations), as well as median values from each experiment (large dots) are shown. Scale bar: 50 µm. Paired t-test.

**Figure 3.**
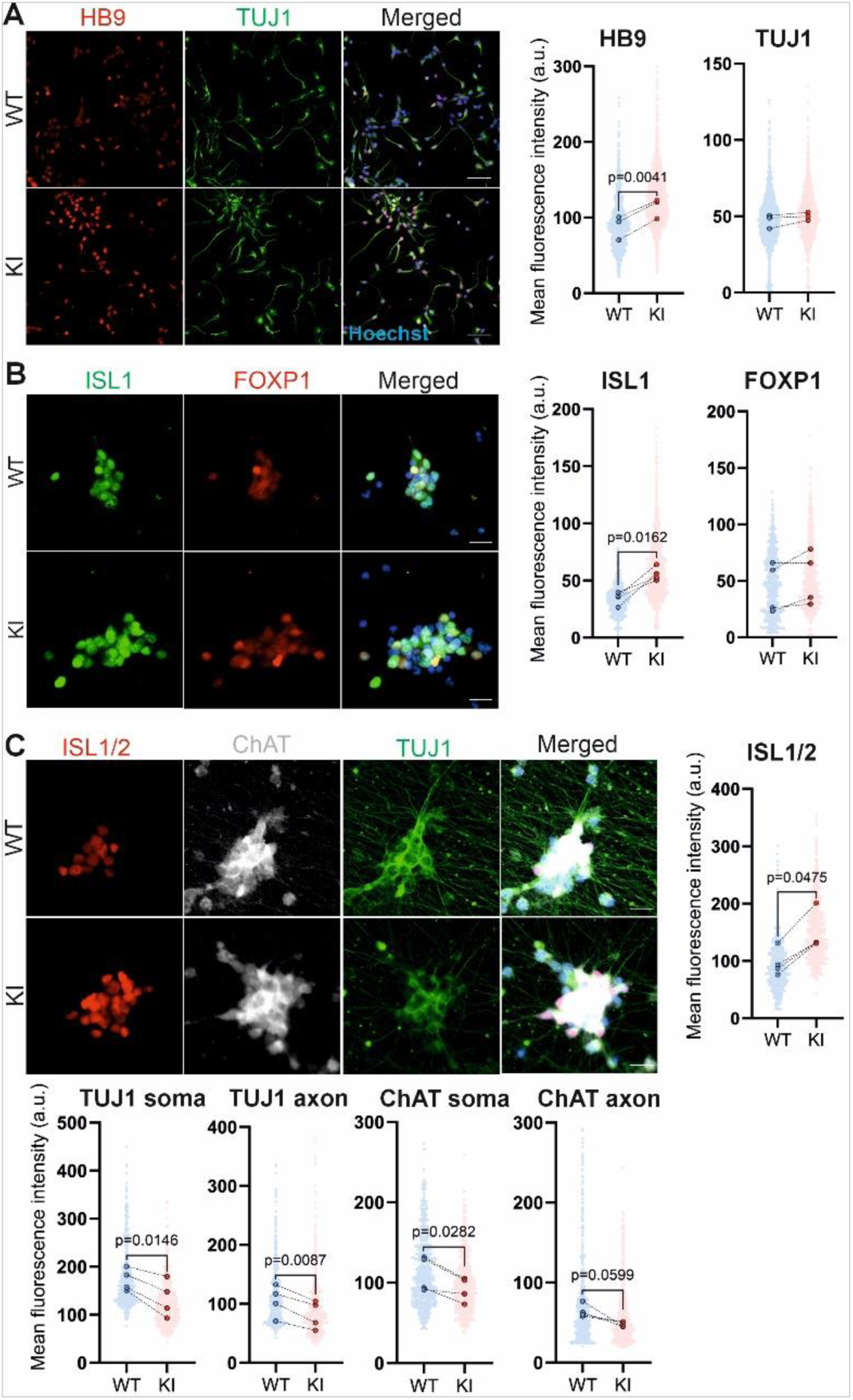
Differentiation of control and LCCS1 spinal motor neurons. **A)** Immunostaining of postmitotic motor neuron progenitors at D11 of differentiation. Spinal motor neuron-specific transcription factor HB9 (red) and an early, pan-neuronal marker TUJ1 (green) are shown in wild type (WT and LCCS1 knockin (KI) differentiated cells. Quantification of mean fluorescent intensity is shown as individual data points generated from each cell (500 cells per differentiation, three independent differentiations), as well as median values from each experiment (large dots). Scale bar: 50 µm. Paired t-test. **B)** Immunostaining of mature motor neurons at D32 for transcription factors ISL1 (green) and FOXP1 (red). Individual data points generated from each cell are plotted (200 cells per differentiation, four independent differentiations), as well as median values from each experiment (large dots). Scale bar: 20 µm. Paired t-test. **C)** Immunostaining of mature motor neurons at D32 of differentiation protocol with motor neuron specific transcription factors ISL1/2 (red) and choline acetyltransferase ChAT (white) and pan-neuronal TUJ1 (green). Quantification of signals and plotting of individual data points generated from each cell (200 cells per differentiation, four independent differentiations). Median values from each experiment are shown as large dots. Scale bar: 20 µm. Paired t-test.

### Choline acetyltransferase insufficiency and neurofilament alterations suggest functional immaturity of LCCS1 motor neurons

The final maturation of pluripotent stem cell derived motor neurons occurs by D32 in culture (18). TUJ1 is an important determinant of neuronal identity while choline acetyltransferase (ChAT) synthetizes the neurotransmitter acetylcholine at the neuromuscular junctions thus enabling the signal transduction as a command for muscle contractions and other motor behaviors (22). Characterization of motor neuron cultures from WT and LCCS1-hESCs at this point demonstrated comparable signal for FOXP1, which is necessary for motor neuron patterning (23) but also an increased intensity of ISL1 (**Fig. 3B**) Accordingly, analysis of the differentiated motor neurons with ISL1/2 specific antibody verified its increase. Contrarily, immunocytochemistry of LCCS1 motor neurons exposed decreased TUJ1 and ChAT (**Fig. 3C**). Simultaneously with these motor neuron functionality and identity changes, we detected significantly reduced perinuclear and cytoplasmic amounts of GLE1 protein in LCCS1 motor neurons (Fig. S2B). These findings suggest that the LCCS1 variant causes disturbed motor neuron cell specification and maturation processes, affecting particularly the transition to the most functionally mature states.

To further characterize the hESC-derived motor neurons, we next assessed their neurofilament expression. The scaffolding/structural protein subunits neurofilament light (NF-L), medium (NF-M), and heavy (NF-H) contribute to the axonal cytoskeleton of myelinated axons and support their cell-type specific size and shapes (24). Analysis of different neurofilament subunits by Western blotting showed comparable levels of NF-L, -M, and -H in WT and LCCS1 motor neurons (**Fig. 4A**). Intriguingly, using immunofluorescent staining we identified two-fold increase in NF-L inclusion incidence in LCCS1 motor neurons (**Fig. 4B**). This data demonstrates that while we initially found only minor motor neuron identity changes, the LCCS1 motor neurons exhibit a disease-relevant pathology characterized by abnormal neurofilament accumulation (25, 26).

**Figure 4:**
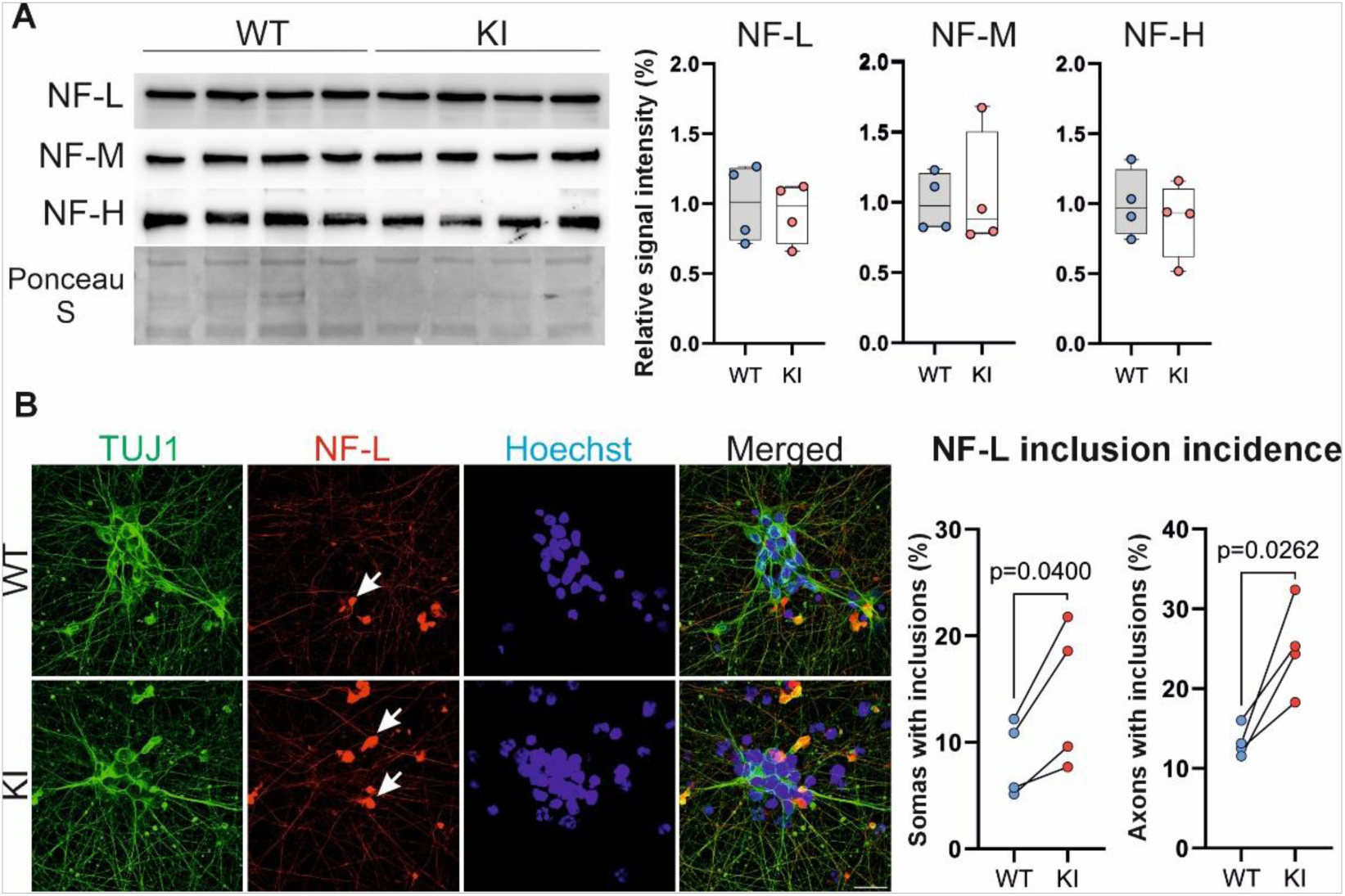
Neurofilament turnover alteration in mature LCCS1 motoneurons indicates functional deficits. **A)** Analysis of neurofilament (NF) Light (L)/Medium (M)/Heavy (H) subunits by Western blot in wild type (WT) and knockin (KI) mature motor neurons (MNs). Quantification data are presented as box plot and whiskers, including individual data points from each differentiation (four independent differentiations). **B)** Visualization and quantification of neurofilament-light (NF-L, red) inclusions (white arrows) in the soma/axon of mature MNs, co-stained with TUJ1 (green). Mean inclusion incidence (%) presented from each differentiation (over 170 cells per differentiation analyzed from four independent differentiations). Scale bar: 20 µm. Paired t-test.

### Transcriptional profiling of the LCCS1 gastruloids identifies alterations in neuromuscular and neural crest derivative development

Recent rapid development of pluripotent stem cell-based *in vitro* models provides a powerful tool to examine human embryogenesis, lineage specification, and the first steps of neurulation in human embryos (27). We assessed the developmental competence of WT and LCCS1-hESCs by their *in vitro* differentiation into gastruloids (28). These artificial posterior embryo-like morphological structures include a neural tube flanked by segmented somites and diverse other cell types, such as neural crest, neural progenitors, renal progenitors, and myocytes (28, 29). LCCS1-hESC-derived gastruloid differentiation did not quantitatively or temporally differ from gastruloids derived from WT hESCs (**Fig. 5A**). Principal component analysis from transcriptomic profiling showed greater variation in LCCS1-derived gastruloids compared to WT (**Fig. S4A**). We identified total of 114 differentially expressed genes (DEGs, log2fold change |1|; adjusted P-value < 0.05) in LCCS1-derived gastruloids (**Fig. 5B**, **Table 1**). Of these, 30 were downregulated and 84 were upregulated transcripts (**Fig. 5B, S4B**). DEGs included remarkably high number of human specific zinc finger (ZNF) transcripts (three down- and 14 upregulated) with predicted functions in the regulation of transcription by RNA polymerase II (https://www.genecards.org/). Notably, we found a significant downregulation of *GLE1* itself in LCCS1 gastruloids, suggesting that the c.432-10A>G variant leads to haploinsufficiency during early differentiation processes.

**Figure 5.**
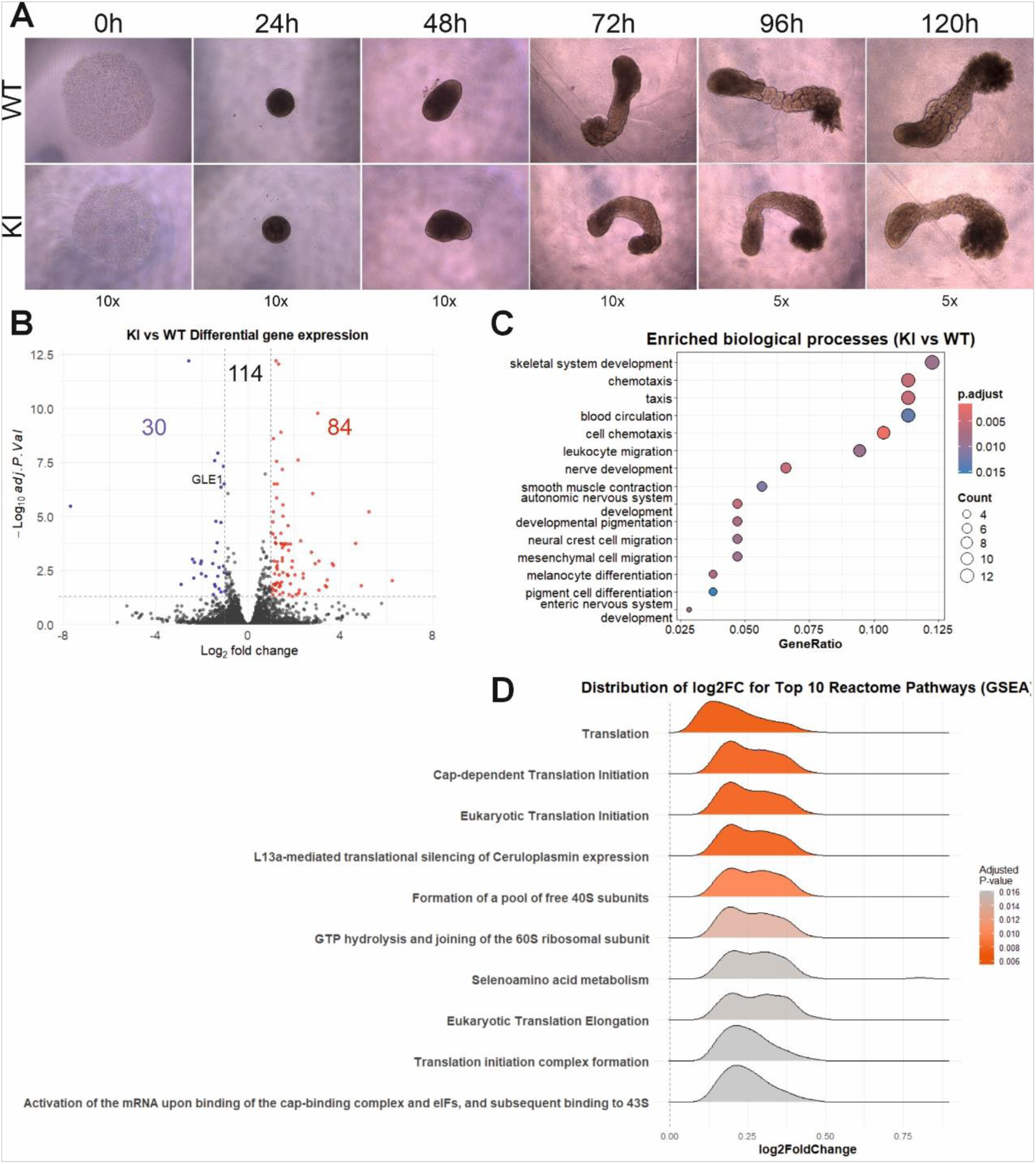
Gastruloid differentiation and transcriptional profiling. **A)** Light microscopy documentation of stepwise differentiation human gastruloids derived from wild type (WT) and LCCS1 knockin (KI) embryonic stem cells. **B)** Volcano plot showing transcriptional changes in LCCS1 gastruloids (114 differentially expressed genes, 30 downregulated, 84 upregulated (log2fold change |1|; adjusted P-value < 0.05). **C)** Gene Ontology pathway analysis was used to identify the enriched biological processes among the DEGs with log2 fold change |1|; adjusted P-value < 0.05. The size of the circle indicates gene counts representing the given biological process and color shows its significance. **D)** Reactome pathway analysis of log2 fold change (log2FC) distributions among differentially expressed genes with the gene set enrichment analysis (GSEA). The color of the distribution plot indicates the significance of the identified cellular pathway.

Pathway analysis of all DEGs revealed significant changes in skeletal system development, chemotaxis, nerve and autonomic nervous system development, pigmentation, neural crest migration and enteric nervous system development. Reactome analysis demonstrated alterations in different aspects of translation as well as ribosome biology (**Fig. 5C-D**). These results demonstrate the capability of LCCS1 hESCs to differentiate into gastruloids but present deficiencies in neuronal differentiation, supporting the previous findings in the human pathology (2). Interestingly, our transcriptomic analyses of LCCS1 gastruloids additionally suggest defects in neural crest development, which were present in the KI mouse model of the disorder (15) but not previously linked to LCCS1 pathology. Collectively, these findings suggest that LCCS1 may represent a neurocristopathy with developmental consequences beyond the motor neuron deficits.

### LCCS1 hESCs-derived ectodermal organoids and LCCS1 fetuses show defects in neural crest differentiation

Human LCCS1 fetuses display deformed skull, small jaw, and pale skin (2), while *Gle1* KI mouse fetuses carrying the PFQ amino acid insertion show changes in the volume of paravertebral sympathetic ganglia, number of adrenal gland chromaffin cells, and branching morphogenesis of sympathetic innervation in the developing heart (15). These features could arise from deficiencies in neural crest derivative differentiation (30, 31). To specifically address this hypothesis, we generated ectodermal organoids from hESCs, allowing the cells to form spheroids from which neural crest cells migrate out (32, 33). This ectodermal organoid differentiation yields early developing ectodermal epithelium both during the neural crest specification and during its early migratory phase.

We observed that both WT and LCCS1 hESCs-derived readily formed ectodermal organoids after exposure to neural crest differentiation medium. Both organoids spontaneously adhered to the bottom of the culture chamber and migratory mats of cells emerged from the organoids (n = 25 organoids/genotype, three experimental replicates). To assess whether the migratory cells in the LCCS1 hESCs-derived organoids are differentiated neural crest cells, we immunostained them with SOX9 antibody, a neural crest specification marker, and SOX2, a marker of the neural plate and neural tube cells (34). We observed SOX2 staining in the adhered organoid and the immediate surrounding whereas the migratory cell mat was negative in both WT and LCCS1 hESCs-derived samples (**Fig. 6A**). SOX9 staining was detected throughout the samples from the adhered organoid to migratory cell mats in WT and LCCS1 hESCs-derived samples (**Fig. 6A**). Using the mean SOX9/SOX2 signal intensity ratio as a proxy of neural crest population we quantified it in organoids, organoid edges, and migratory cells (**Fig. 6B**). Although this did not reveal statistically significant differences between groups a consistent trend toward a reduced ratio was observed in LCCS1 hESC-derived organoids. Consistent with this, we did not observe statistically significant decrease in the mean intensity of the SOX9 in the migratory cells of WT or LCCS1 hESC-derived organoids (**Fig. 6C**). To assess the purity of the migratory neural crest cells, we quantified on average 99.13% of SOX9+ and 1.06% of SOX2+ cells in the migratory cells of the WT samples (**Fig. S4C**). Similarly, an average of 98.84% LCCS1 hESC-derived migratory cells were SOX9+ and 0.34 % were SOX2+ (**Fig. S4C**) indicating significant enrichment of the migrating neural crest cells in both genotypes but showing slightly diminished yet statistically not significant proportions in the LCCS1 model.

**Figure 6.**
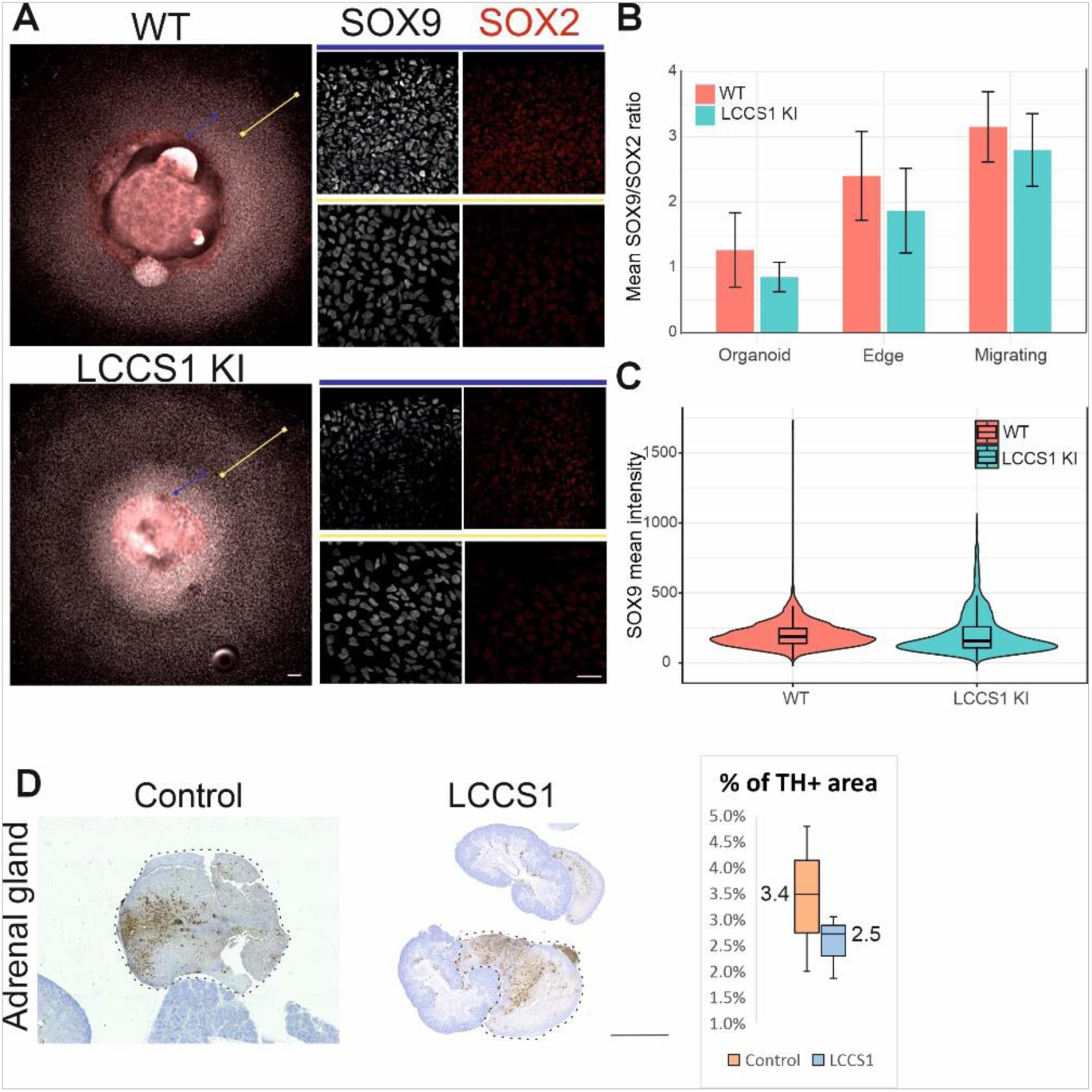
Neuruloids derived from LCCS1 hESCs and neural crest differentiation in LCCS1 fetuses. **A)** Representative images of ectodermal organoids from wildtype (WT) and LCCS1 knockin (KI) cells immunolabelled against SOX9 (white) and SOX2 (red). Representative high-resolution images acquired from organoid edge migrating neural crest cells. Blue lines depict sphere edges; yellow lines portray migrating neural crest cells. Scale bars: 200µm and 20µm **B)** Mean SOX9/SOX2 ratio of organoid, organoid edge, and migrating neural crest nuclei from WT and LCCS1-KI organoid samples. No statistical difference was observed between the organoid (WT 1.27±0.57, n=4 vs LCCS1-KI 0.85±0.22, n=4), organoid edge (WT 2.40±0.68, n=7 samples vs. LCCS1-KI 1.87±0.65, n=5 samples), or migrating neural crest cell (WT 3.15±0.54, n=5 samples vs. LCCS1-KI 2.80±0.56, n=5 samples) in Wilcoxon rank sum test. Column and whiskers represent mean and standard deviation of the mean SOX9/SOX2 ratio in each sample. Black dots represent the mean Sox9/Sox2 intensity ratios of different samples. **C)** Mean SOX9 intensity in nuclei of migrating neural crest cells from WT and LCCS1-KI. No significant difference was measured between WT (202±89 a.u., n=3516 cells from 2 independent experiments) and LCCS1-KI (207±147 a.u., n=1719 nuclei from 2 independent experiments) samples (P=0.211 Wilcoxon rank sum test with continuity correction). **D)** Representative cross-sections of human control (H13+3) and LCCS1 (H14) adrenal glands, which are circled by dotted lines and immunostained for tyrosine hydroxylase (TH) representing the chromaffin cells. Quantification of mean TH-positive area (± SD) within the adrenal glands is shown (*n* = 3 per genotype, 3-8 sections per fetus). Scale bar 2000 μm.

Ectodermal organoid results indicate that the primary neural crest cell specification and early migration are not significantly affected in LCCS1 hESC-derived organoids but also suggest consistent minor variations that may alter downstream differentiation in target organs (35, 36). We used retrospective pathology samples collected from three LCCS1 and three control fetuses aged from gestational week 13+2(H13+2) to 20+3 (**Table 2**) to address neural crest contribution in adrenal gland differentiation. The developing adrenal gland is proportionally larger than the mature organ (37) and it is of mesodermal origin except for the medullar chromaffin cells, which originate from neural crest and migrate there during the second (38). Thus, fetal adrenal glands were used as a proxy of neural crest derivative development in LCCS1 patients. Quantification of tyrosine hydroxylase (TH)-positive chromaffin cell density, normalized to total adrenal gland area, demonstrated a significant reduction in chromaffin cells in LCCS1 adrenal glands compared with age-matched controls (**Fig. 6C**, 3.4% ± 0.022 in control vs. 2.5% ± 0.017 in LCCS1, Welch’s t-test p = 0.0385). This suggests a defect in neural crest biology and is in line with our previous studies hypothesizing LCCS1 as a neurocristopathy.

### LCCS1 motor neurons and patient-derived fetal fibroblasts accumulate RNA despite substantially lower transcription and translation activity

Previous studies addressed the cellular consequences of *GLE1* c.432-10A>G variant in yeast and HeLa cells by overexpressing mutant GLE1 and identified remarkable accumulation of poly(A)+ RNA in the nucleus (11) However, our work in mouse cells and tissues originating from the LCCS1 KI mice did not find abnormalities in poly(A)+ RNA distribution (15). To examine this phenotype in hESCs and hESC-derived motor neurons, we performed fluorescent *in situ* hybridization (FISH) with an oligo d(T) probe. This demonstrated slightly increased general poly(A)+ RNA signal in undifferentiated LCCS1-hESCs but not in the postmitotic motor neuron progenitors at D11 (**Fig. S5A-B**). D32 mature motor neurons derived from LCCS1-hESCs showed homogeneously increased oligo d(T) intensity in the nucleus and cytoplasm, but leaving the nucleocytoplasmic distribution intact in LCCS1-hESC-derived motor neurons (**Fig. 7A**).

**Figure 7.**
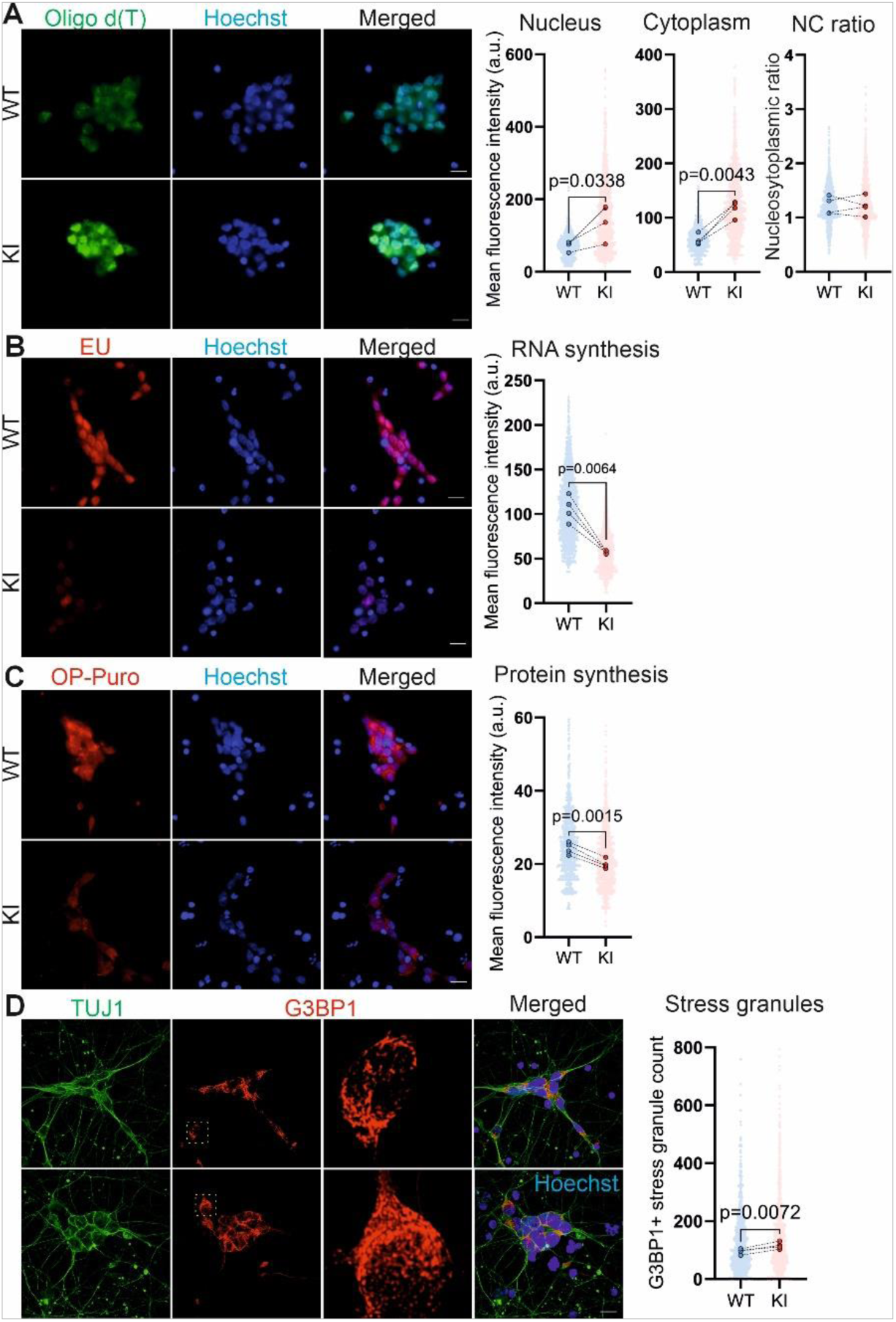
GLE1-mediated cellular functions in mature motor neurons carrying LCCS1 variant. **A)** Oligo d(T) probe fluorescent *in situ* hybridization (FISH) to detect the poly(A)+ RNA and quantify its subcellular abundance and nucleocytoplasmic distribution in mature wild type (WT) and LCCS1 knockin (KI) motor neurons (MNs). Hoechst visualizes all nuclei. Individual data points generated from each cell are plotted (400 cells per differentiation, three independent differentiations), as well as median values from each experiment (large dots) are shown. Scale bar: 20 µm. Paired t-test. **B)** Quantification of RNA synthesis in mature wild type (WT) and LCCS1 knockin (KI) MNs. Ethynyl uridine (EU, red) incorporation (1mM EU, 1 hour pulse) together with Hoechst visualizing all nuclei are shown. Individual data-points generated from each cell are plotted (400 cells per differentiation, four independent differentiations), as well as median values from each differentiation (large dots). Scale bar: 20 µm. Paired t-test. **C)** Protein synthesis in mature wild type (WT) and LCCS1 knockin (KI) MNs by O-propargyl-puromycin (OP-Puro, red) metabolic labelling (20 µM OP-Puro, 30 min pulse). Hoechst visualizes all nuclei. Individual data points generated from each cell are plotted (250 cells per differentiation, four independent differentiations), as well as median values from each differentiation (large dots) shown. Scale bar: 20 µm. Paired t-test. **D)** Induction of hyperosmotic shock (400 mM sorbitol, 60 min) and quantification of G3BP1-positive (red, zoomed in areas in yellow squares) stress granule in mature MNs visualized by TUJ1 (green) immunostaining. Individual data points generated from each cell are plotted (400 cells per differentiation, three independent differentiations), as well as median values from each experiment (large dots) shown. Scale bar: 20 µm. Paired t-test.

To determine whether the increased steady-state RNA levels result from enhanced transcription, we quantified the nascent RNA synthesis using 5-ethynyluridine (EU) metabolic labeling. Contrary to this hypothesis, LCCS1 motor neurons exhibited significantly reduced transcription compared with WT controls (**Fig. 7B**). Consistent with these results, protein synthesis measured by incorporation of O-propargyl-puromycin (OP-Puro) remained unchanged in undifferentiated hESCs and postmitotic motor neuron progenitors but was significantly diminished in LCCS1-hESC-derived motor neurons (**Fig. 7C, S5C-D**). Notably, the transcriptional and translational changes were accompanied by a pronounced increase in cellular stress responses. LCCS1-hESC-derived motor neurons displayed a significant increase in G3BP1-positive stress granules following 60 min of hyperosmotic stress (Fig. 7D), closely mirroring the phenotype previously reported following siRNA-mediated GLE1 depletion in HeLa cells (10). These findings highlight stress-granule dysregulation as a robust and conserved consequence of impaired GLE1 function. To assess whether altered translation reflected changes in ribosome engagement, we performed polysome profiling followed by qPCR in mature motor neurons. We did not detect significant differences in polysome association of the analyzed transcripts between genotypes (**Fig. S6A-B**), suggesting that translational repression in LCCS1-hESC-derived motor neurons correlates with stress-granule accumulation rather than to changes in ribosome loading.

A reduction in protein output can reflect either impaired synthesis or enhanced degradation. To test potential changes in protein degradation, we examined two main protein clearance pathways: ubiquitin-mediated proteasomal degradation and autophagy. Neither total protein ubiquitination nor the abundance of the autophagy markers p62 and LC3 differed between control and LCCS1 motor neurons (**Fig. S6C-D**). This indicates that global protein degradation does not increase, and that the reduced protein synthesis in hESC-derived motor neurons does not arise from enhanced protein clearance.

Next, we analyzed RNA distribution, protein synthesis and stress reactions in human patient-derived fetal fibroblasts. These identified similar cellular responses as detected in the LCCS1-hESC-derived motor neurons. While poly(A)+ RNA distribution and abundance was unaffected, global protein translation was significantly diminished, and the G3BP1-positive stress granule count were higher in hyperosmotically shocked patient cells (**Fig. 8A-C**). In summary, contrary to the originally proposed pathogenic mechanism of LCCS1, we found that the effects of LCCS1 variant on GLE1 function depend on the cell’s differentiation status and identity rather than on RNA distribution.

**Figure 8.**
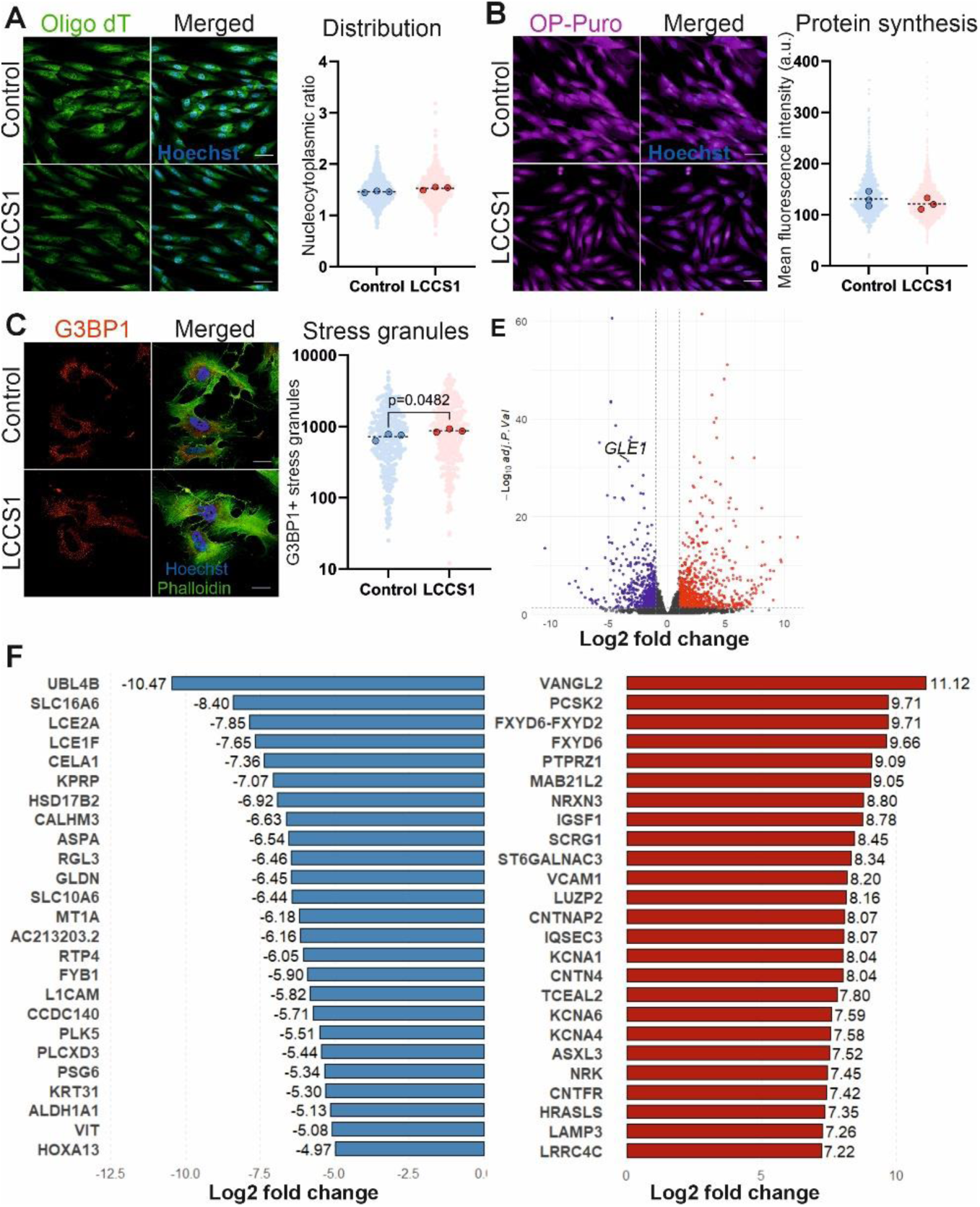
GLE1-mediated cellular functions in LCCS1 fetus-derived fibroblasts. **A)** Visualization and quantification of poly(A) + RNA nucleocytoplasmic distribution by fluorescent *in situ* hybridization of 5′-Cy3-oligo d(T)30 probe (green) in control and LCCS1 fibroblasts. Cell nuclei are visualized by Hoechst staining (blue). Individual data points generated from each cell are plotted (300 cells per patient, three patients per genotype) and median values (large dots) from each patient are presented. Scale bar 50 μm. **B)** Quantification of protein synthesis by metabolic incorporation of O-propargyl-puromycin (OP-puro) (20 μm OP-Puro, 30 min pulse) in control and LCCS1 fibroblasts. Individual data points generated from each cell are plotted (500 cells per patient, three patients per genotype) and median values (large dots) from each patient are presented. Scale bar 50 μm. **C)** Induction of hyperosmotic shock (400 mM sorbitol, 60 min) and visualization of stress granule assembly factor 1 (G3BP1)-positive (red) stress granules in the cytoplasm (phalloidin, green) of control and LCCS1 fibroblasts. Stress granule quantity shown as individual data points of total stress granule count detected in each cell (100 cells per patient, three patients per genotype). Median values (large dots) from each patient are also demonstrated. Scale bar 30 μm, two-tailed Student’s t-test. **D)** Volcano plot showing the 1343 differentially expressed genes (DEGs) in bulk RNA-seq analysis of LCCS1 fibroblasts in comparison to healthy controls (three biological replicates per genotype). Significantly downregulated genes in LCCS1 fibroblasts are indicated in blue and upregulated red (log_2_ fold change |1|; adjusted *P*-value < 0.05).

### LCCS1 cells show changes in mRNA decay rate

Our results in LCCS1-hESC-derived motor neurons and patient fetal fibroblasts (**Figs. 7-8**) indicate that abnormal poly (A)+ RNA localization or RNA accumulation cannot explain deficits in LCCS1 cellular homeostasis. To analyze LCCS1 cells in an unbiased manner and test whether altered RNA decay contributes to pathology, we conducted global transcriptomic profiling in intact and transcriptionally inhibited (D-actinomycin) control and patient cells, which both clustered among their own groups (Fig. S7A-B). We detected a total of 1343 DEGs (log2fold change |1|; adjusted P-value < 0.05) in intact patient cells, of which 795 were upregulated and 548 downregulated (**Fig. 8E-F**. Ubiquitin Like 4B (*UBL4B*), operating within the complex ubiquitin system, was the most downregulated transcript in LCCS1 cells (Fig. 8F). In general, downregulated transcripts related to skin identity (*LCE2A*, *LCE1F*, *KPRP*), transmembrane transport activity (*SLC16A6*, *CALHM3*, *SLC10A6*). Interestingly, this list also included gliomedin (*GLDN*), the causative gene for LCCS11, which is also a lethal congenital arthrogryposis (39, 40). Of the top upregulated transcript VANGL planar cell polarity 2 (*VANGL2*) and Proprotein Convertase Subtilisin/Kexin Type 2 (*PCSK2*), the mutations of which contribute to neural tube defects as well as myocardial infarction, respectively (41, 42). Other upregulated genes play roles in cell adhesion (*IGSF1*, *VCAM1*), ion transport regulation (*FXYD6*, -*2*) and potassium voltage-gated channel (*KCNA1*, -*4*, -*6*) (**Fig. 8F**).

In patient-derived fetal fibroblasts, global transcriptome analysis after 1, 3, and 9 hours of transcriptional inhibition via D-actinomycin (10mg/ml) identified 367 transcripts with reliable half- life estimates (R² ≥ 0.5) in at least one condition (either Control or LCCS1 cells) (**Fig. 9A, Table 4**). Overall, the distributions largely overlapped, indicating that overall RNA stability is not profoundly altered in LCCS1 fibroblasts. Consistent with this, the vast majority of transcripts showed no significant difference in decay rate between genotypes. Among the 367 transcripts analyzed, only a small subset exhibited altered stability: 34 transcripts showed significantly faster decay and 4 showed increased stability in LCCS1 cells (**Fig. 9B**). For these differentially decaying transcripts, fitted degradation rates (beta values) were modestly but consistently higher in LCCS1 fibroblasts, indicating slightly accelerated RNA turnover (**Fig. 9C**). Transcripts with reduced stability in LCCS1 include ribosomal, ubiquitin, and translation related proteins with disease relevance to anauxetic dysplasia (*POP7*) displaying abnormalities in skeleton, joints, and facial features, developmental and epileptic encephalopathy (*ZNF622*), and early-onset Parkinson disease (*MUL1*). Importantly, GO pathway analysis of the transcripts with significant differences in their degradations rates between LCCS1 and control fibroblasts identified ribosome biogenesis and assembly, protein destabilization, and mitochondrial ribosome assembly as the most affected biological processes (**Fig, 9D**). This lays in line with our polysome profiling results, supporting aberrations in ribosome biology as the cause for mRNA accumulation and subsequent deficits in protein homeostasis in LCCS1 cells.

**Figure 9.**
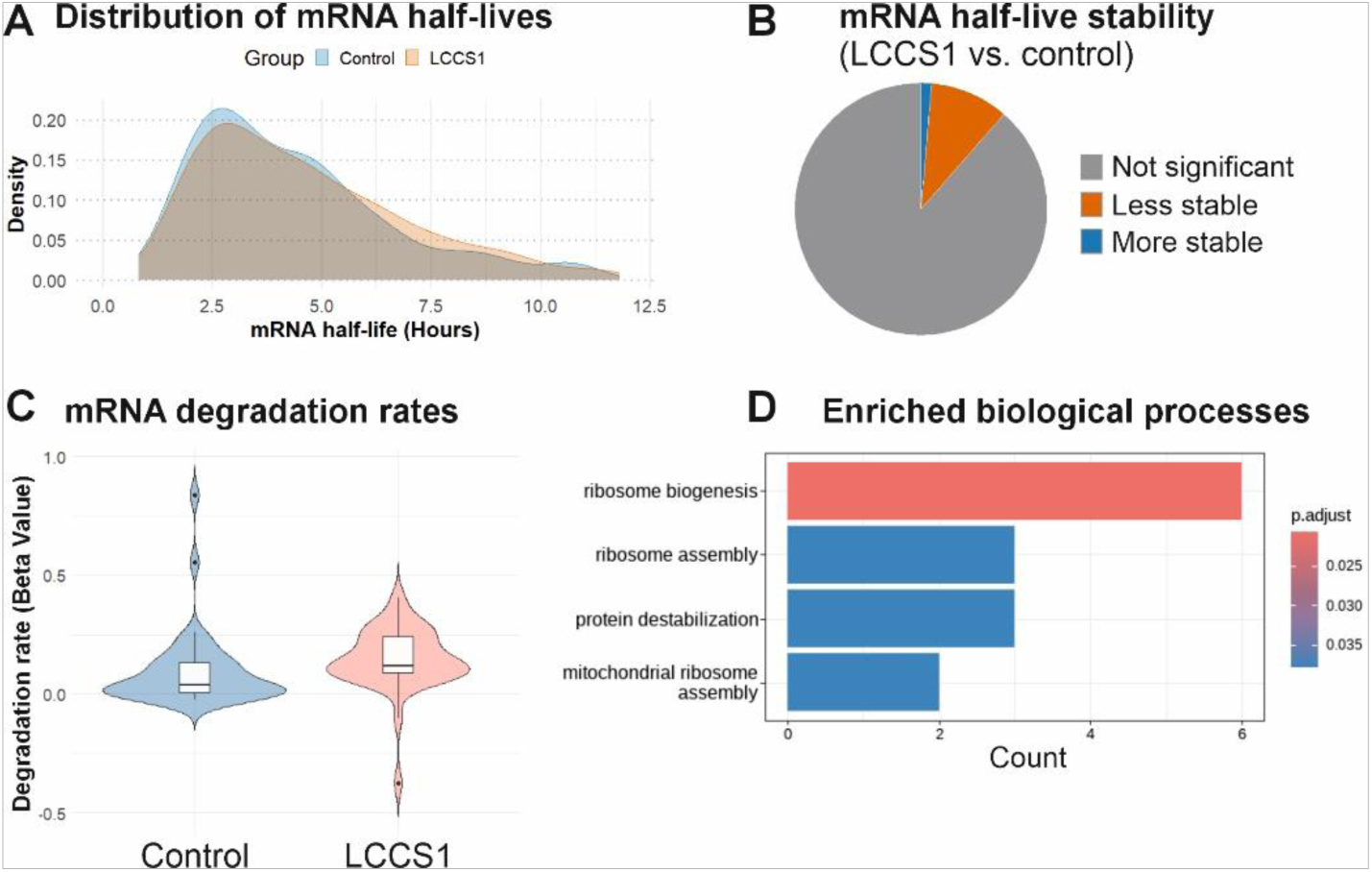
Assessment of mRNA degradation rate in control and LCCS1 fibroblasts. **A)** The density plot displays the distribution of mRNA half-lives in the control (315 transcripts plotted) and LCCS1 (347 transcripts plotted) fetal fibroblasts. The plot includes a total of 367 unique transcripts that had a reliable half-life estimate (R-squared ≥ 0.5) in at least one of the timepoints. The height of each curve indicates the frequency of genes with a specific half-life, allowing for a visual comparison of the distributions. **B)** Pie plot presenting the distribution of mRNA transcripts with higher (blue, 4 out of 367) and lower (orange, 34 out of 367) stability in LCCS1. **C)** Box plot with whiskers presenting the distribution of fitted beta degradation rates (the slope of the fitted exponential decay curve describing mRNA abundance over time) for total 38 transcripts with significant decay differences between LCCS1 and control patient cells. **D)** Gene ontology enrichment analysis of biological processes augmented in LCCS1 fetal fibroblasts. Visualization shows top functions that are related to the 38 transcripts whose degradations rate is significantly different between LCCS1 and Control cells.

### Reduced Mucin-Type O-Glycosylation in LCCS1 Fibroblasts

To assess biological processes and pathways affected by *GLE1*_FinMajor_ variant, we applied GO and Reactome pathway pipelines to DEGs identified by transcriptomic profiling of LCCS1 patient fibroblasts (**Fig. 8E-F**). The most significantly enriched GO biological processes in LCCS1 cells include circulatory system, skeletal system development, and axon development (**Fig. 10A**), along with additional axon-related processes. Reactome pathway analysis revealed changes in neuronal system and many electrical transmission-related pathways, highlighting changes in neurobiological functions in LCCS1 cells (**Fig. 10B**).

**Figure 10.**
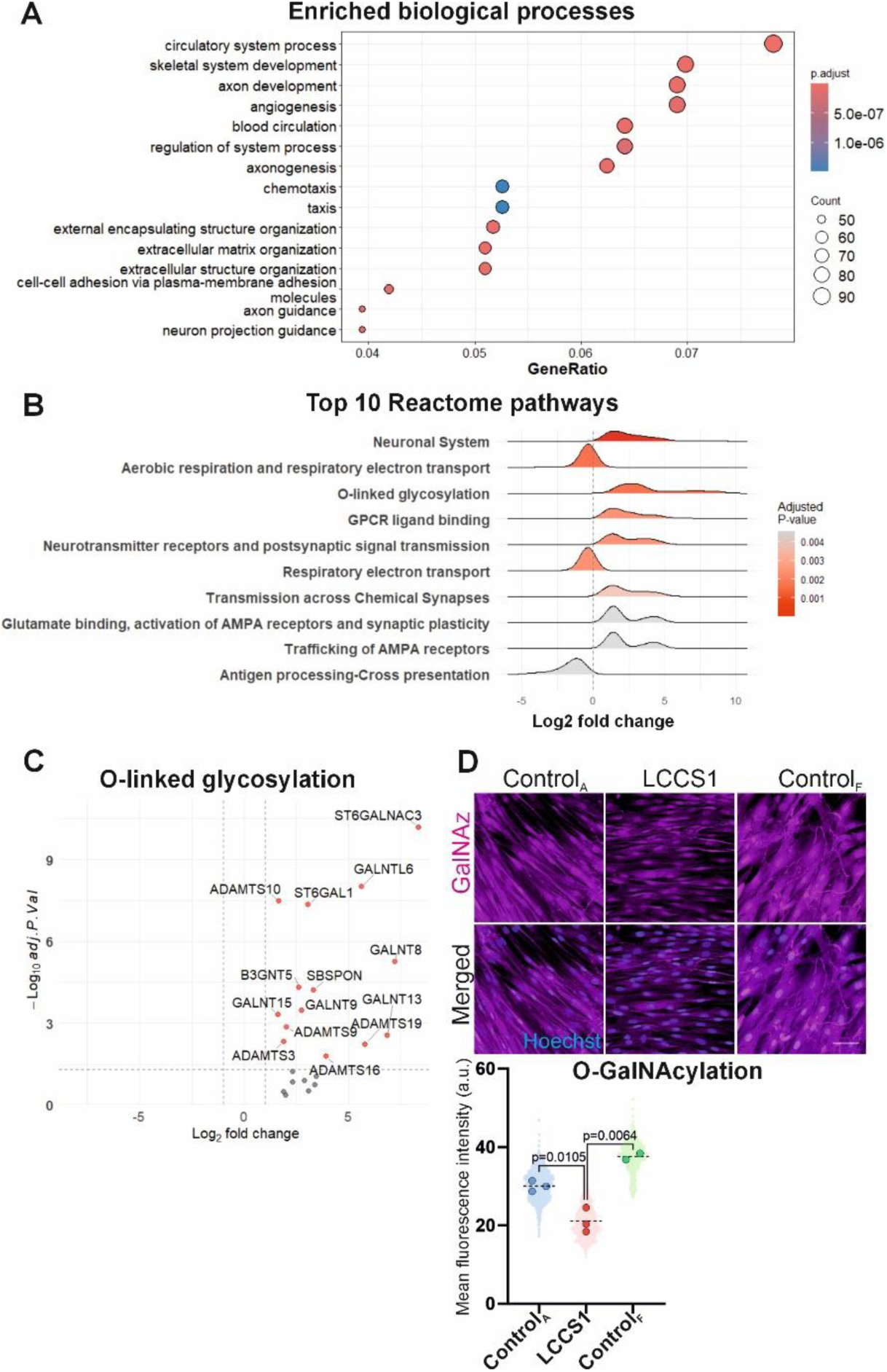
Transcriptional profiling of LCCS1 fibroblasts identifies major changes in protein post-translational modifications. **A)** Dot plot illustrates the gene ontology enrichment analysis of the top 15 molecular functions impacted by the *GLE1*_FinMajor_ variant. Transcripts with adjusted *P*-value < 0.05 and log_2_ fold change > 1 in either direction were used for analysis. **B)** Ridge plot of functional enrichment analysis displaying the top 10 Reactome pathway terms. **C)** Plot showing a subset of differentially expressed genes linked to the ’O-linked glycosylation’ Reactome pathway. **D)** Metabolic labelling of patient fibroblasts with 50 µM GalNAz (magenta) for 72 hours followed by the click chemistry to quantify the level of O-GalNAcylation in control and LCCS1 derived fibroblasts. Individual data points generated from each cell are plotted (500 cells per patient, three adult control cell lines, 3 LCCS1 patient cell lines, 2 fetal control cell lines), as well as median values (large dots) from each sample are shown. Scale bar: 50 μm.

Interestingly, Reactome analysis also identified a rare non-neuronal pathway, O-linked glycosylation affected in LCCS1 fibroblasts (**Fig. 10B)**. Glycosylation is known as a critical post-translational regulator during development and cancer (43, 44). The identified transcripts represent several N-acetylgalactosaminyltransferase (*GALNT* 6, 8, 9, 13 and 15) enzymes (**Fig. 10C**). These proteins initiate non-mucin type O- linked N-acetylglucosaminylation (O-GlcNAcetylation), the addition of N-acetylglucosamine (GlcNAc) to the substrate proteins. They also act in mucin type N-acetylgalactosaminylation (O-GalNAcylation), which adds N-acetylgalactosamine (GalNAc) to proteins. While changes in the O-GalNAcylation are mainly reported in the context of cancer (45), O-GlcNAcylation affects mammalian neurodevelopment (46).

To assess whether the identified transcriptional changes related to post-translational modifications have functional consequences, we first analyzed O-GalNAcylation in LCCS1 patient and control cells. We detected a significantly decreased addition of GalNAc monosaccharides in LCCS1 cells (**Fig. 10D**). As O-GalNAcylation typically occurs in the Golgi apparatus (45), we immunostained LCCS1 and control cells with GM100 and analyzed the signal area as a proxy of organelle volume. We found no differences in median Golgi volumes between the cell types (**Fig. S7D**). Lastly, we analyzed O-GlcNAcylation in LCCS1 patient and control cells without detecting significant differences (**Fig. S7C**), indicating that the deficits in O-linked glycosylation remain specific to mucin-type O-GalNAcylation.

## DISCUSSION

We examined cellular and physiological functions of human GLE1 in the devastating fetal-lethal rare neurodevelopmental disorder LCCS1. The condition manifests already *in utero* and inevitably leads to death of the fetus before the 32^nd^ gestational week (2). While the cause of death in human LCCS1 patients remains elusive, our recent study with LCCS1 KI mouse model points towards deficits in adrenal gland medulla, paravertebral sympathetic ganglia, and cardiac sympathetic innervation (15). The mice present similar but clearly milder motor neuron and spinal cord phenotype spectrum as the human patients, either due to species specific alterations in splicing of *Gle1*/*GLE1* or differences in physiological functions of RNA metabolism.

Previous studies of GLE1 function and binding partners have established GLE1 as a multifunctional DEAD-box protein RNA helicase regulator, that contributes to quality control throughout the mRNA life cycle (9–12). The first experiments addressing the functional consequences of the *GLE1* c.432-10A>G variant overexpressed a tagged pathogenic GLE1 protein *in vitro* in HeLa and yeast cells (11). These studies suggested that disturbed mRNA export results in nuclear mRNA accumulation in LCCS1 cells and causes cellular defect of the LCCS1 pathology. Interestingly, knockdown of *GLE1* in HeLa cells not only impairs RNA transport, but also reduces general nascent protein translation and increases the stress granule count in cells experiencing challenging conditions (10).

To gain physiologically and clinically relevant insights into RNA biology underlying LCCS1-associated neurodevelopmental and degenerative disorders, we combined studies of hESC-derived differentiated cells with patient-derived cells and tissues. Our studies revealed that *GLE1* c.432-10A>G variant does not interfere with nucleocytoplasmic poly(A)+ RNA distribution in human cells despite the clearly reduced levels of mutant GLE1 protein. The variant has cell-type specific effects on cellular propagation as detected by diminished proliferation in LCCS1 hESCs and increased proliferation in patient fibroblasts. Furthermore, we detected consistent decrease in global transcription as well as in translation, changes in mRNA decay kinetics and increased stress granule formation specifically in mature cell type. These cellular findings largely align with our recent mouse studies. There, we showed that the nine-nucleotide insertion mimicking the alternative splicing of *GLE1* c.432-10A>G variant does not affect the nucleocytoplasmic distribution of poly(A)+ mRNA in mouse primary cells or embryonic day 11.5 (E11.5) brachial spinal cords (15). Instead, the *Gle1* KI fetal primary cells presented decreased proliferation and disturbed cell cycle progression, as well as significantly diminished overall RNA transcription and protein translation. Thus, our studies strongly imply that the c.432-10A>G variant severely affects GLE1A isoform associated functions, translation regulation and stress granule formation (10, 47, 48).

Profiling of mRNA half-lives in control and LSCC1 patient-derived fetal fibroblast identified ribosome biogenesis and assembly as well as of protein destabilization as affected biological processes in LSCC1. Although we cannot exclude the possibility that diminished translation derives from diminished overall RNA synthesis in LCCS1 cells, our group recently detected diminished translation, formation of G3BP1-positive stress granules and significant dysregulation of micro-RNA expression in LCCS1 KI mouse primary cells exposed to stressful conditions (15). Interestingly, DDX3, a DEAD-box RNA helicase and previously reported interaction partner of GLE1, plays a crucial role in ribosome assembly, specifically by promoting the formation of functional 80S ribosomes and translation initiation (49). DDX3 also regulates the biogenesis of certain cancer-related micro-RNAs, which provides/suggests an interesting mechanistic link for defects in protein synthesis in LCCS1 cells, together with changes in RNA decay rates identified in this study.

Transcriptomic profiling of fetal LCCS1 patient-derived fibroblasts identified O-linked glycosylation, specifically very little studied O-GalNAcylation, and protein destabilization as affected post-translational changes. Post-translational modifications play a key role in translation regulation, and among them, ubiquitin-proteasome system becomes increasingly important for determining the correct balance between protein synthesis and degradation (50). Ubiquitination controls ribosomal function and quality during translation, and issues in these processes can cause neurodevelopmental disorders (51). Recent research highlights GLE1 association with ubiquitin-proteasome system via interactions with E3 ubiquitin ligases makorin 2 (MKRN2), TDP-43 and Hsp90 (12, 52). Increased interaction with DDX1 and GLE1A/DDX3-dependent translation stalling occur especially under stressful environment, both resulting in the formation of stress granules and linking cellular stress with ubiquitin-mediated signaling (4, 10, 47, 53). We detected significant increased NF-L inclusion incidence in LCCS1 hESC-derived motor neurons highlighting disease-relevant pathologically abnormal neurofilament accumulation and suggesting changes in ubiquitin ligase functions (54, 55). Increased expression of NF-L produces morphological alterations in murine motor neurons while NF-L inclusions typically occur in human motor neuron disease pathology (25, 26).

Aberrant RNA handling and translational control are key attributes of cellular stress while RNA toxicity and translation dysregulation are well-known pathologies of neurodegeneration (56–58). Our results in LCCS1 hESC-derived motor neurons and fetal LCCS1 patient-derived fibroblasts indicate that abnormal RNA localization alone cannot explain LCCS1 defects in cellular homeostasis. Patient fibroblasts and hESC-derived motor neurons display normal nucleocytoplasmic poly(A)+ RNA distribution, indicating that the *GLE1* c.432-10A>G variant does not universally cause nuclear RNA overload. This suggests that the cellular dysfunction in LCCS1 arises from impaired handling of RNA under conditions that demand high post-transcriptional control, such as neuronal differentiation and stress.

## METHODS

### hESC culture

The H9 human embryonic stem cells (hPSCReg ID: WAe009A) were cultured on 3.5 cm dishes coated with growth factor reduced Matrigel® (diluted in 1:200 ratio in DMEM/F12 media with GlutaMAX: 10565018, Thermo Fisher) and maintained in Essential 8™ medium (A1517001, Thermo Fisher) in Heraeus Heracell 150 Incubator at 37 °C with 5% CO_2_.

Cells were passaged when they reached 70-80% confluence by mechanically scraping the cells in the presence of 0.5 mM EDTA in PBS. First, the cells were washed twice with EDTA (1ml/well on a six well plate) and then incubated in EDTA 3-4 minutes at 37°C. E8™ + 5 μM ROCKi was used for re-suspension and re-seeding of the cells. A day after passaging, the ROCKi was removed by changing the medium into plain E8™. The cells were monitored daily to ensure their undifferentiated state.

### Fibroblast culture

The patient LCCS1 fibroblasts, as well as healthy fetal and adult controls, were cultured in DMEM medium (31966-021; Gibco, Bleiswijk, Netherlands) + 10% FBS (10500064; Thermo Fisher) + penicillin/streptomycin (15140122; Thermo Fisher, New York, USA). All of the experiments with fibroblasts were performed at passage 4 on glass fibronectin (1030-FN-05M; R&D Systems, Minneapolis, MN, USA) coated coverslips or plastic culture plates (Starlab; Hamburg, Germany).

### Electroporation-based CRISPR/Cas editing

The hESCs were grown to 70-80% confluency and three hours before electroporation, 10 μM ROCKi was added to the cells. To assemble the ribonucleoprotein (RPN) complex before electroporation, 100 pmol of crRNA (Alt-R® A.s. Cas12a crRNA, IDT, 100 µM) were complexed with 75.6 pmol of Cas12a (Cpf1). After incubation, 2 µl of 10.8 μM electroporation enhancer (AltR® Cpf1 Electroporation Enhancer, IDT, 100 µM) were diluted with IDTE buffer and 4 µg of ssDNA repair template was added to the mixture (Supplementary table 1). To dissociate the cells for electroporation, the medium was first removed and then the cells were washed twice with 5% FBS-PBS. Then, the cells were treated in 0.5ml of Accutase™ (A1110501, Thermo Fisher), and incubated at 37°C for 3-4 min. After Accutase™ treatment, the cells were mechanically dissociated into single cells by gently pipetting up and down in Accutase™. The suspension was collected to a 15 mL Falcon tube and the wells were washed twice with 1 ml of 5%FBS-PBS (also collected into the Falcon tube). The cells were centrifuged at 1115 RPM for 3-4 minutes, counted, and about 2 million cells were used per electroporation. The cells were re-suspended to Neon™ resuspension buffer R, and 120 μl of cell suspension was mixed with RNP complex mixture for the electroporation. The electroporation was performed using the Neon™ Transfection system with the 100 µL electroporation tips (Invitrogen™). The cells were electroporated with two 20 ms pulses at 1100 mV. After electroporation, the cells were seeded onto six-well plates containing E8™ medium + 10 μM ROCKi.

### Cell sorting and colony picking

Two days post electroporation, hESCs were scraped after Accutase™ treatment and live single cells were sorted using the Sony SH800Z Cell Sorter (1 cell was plated per one well on a 96- well–plate in 100 µl of E8™ medium containing 10 µM ROCKi, 10% CloneR (#05888, STEMCELL technologies) and 1% Penicillin/Streptomicin (ECB3001, Euroclone)). After single cell sorting, the cells were left to recover undisturbed for 72 h, after which 100 µl of E8™ medium with 5 uM ROCKi was added. The medium was changed every 3-4 days (fresh E8™ with 5 uM ROCKi), until the cells reached high enough confluency (14-21 days). The cells were split by gently scraping the well and dissociating the monoclonal colony. Half of the cells (around 50 000) were taken for genotyping, and the other half were seeded on a 24 -well plate for expansion.

### Genotyping and sequencing

While picking the colonies from the 96-well plates, genotyping samples were collected and lysed with Direct PCR lysis reagent (250-p1bdxp-100, Biosite). The sequence of interest was amplified by PCR reaction via Phusion polymerase mix (F531L, Thermo Fisher), 20 pmol/μl of forward- and reverse primers (IDT, Supplementary table 1), and cell lysate, using the SimpliAmp Thermal Cycler. After PCR, electrophoresis was performed on 2% agarose (BIO41025, Meridian Bioscience) gel in 1x TAE buffer with 12 µl SYBR™ safe DNA gel stain (S33102, Thermo Fisher). The PCR products (573 bp) were cut out from the agarose under the UV-light for DNA purification, using the NucleoSpin PCR clean-up kit (16361616, Fisher scientific). The purified DNA was used for Sanger-sequencing by Eurofins Genomics (Germany) and the electropherogram was viewed in SnapGene software. Upon validation of the correct editing in hESCs, LCCS1-KI cell line was registered to human pluripotent stem cell registry (hPSCReg ID <u>WAe009A-A-90</u>).

### Spinal motor neuron differentiation

To differentiate hESCs into spinal motor neurons (MN), we used previously published protocols (16, 17, 59). MN differentiation base media (MNb) containing DMEM/F12 (11320033, Thermo Fisher), Neurobasal medium (21103049, Thermo Fisher) 1:1, 0.5× N2 supplement (17502048, Thermo fisher), 0.5× B27 (17504044, Thermo Fisher), L-ascorbic acid 0.1 mM (CAS 50-81-7, Santa Cruz) and Primocin 100 μg/ml (ant-pm-05, Invivogen) were used throughout the differentiation. Briefly, on day 0, hESCs were dissociated with 0.5 mM EDTA into small clusters and resuspended into ultralow attachment plates (CLS3471-24EA, Merck) with differentiation media 1; MNb supplemented with 40 μM SB431542 (616464, Merck), 0.2 μM LDN-193189 (SML0559, Merck), 3 µM CHIR99021 (S1263, Selleckchem), and 5 µM Y-27632 (S1049, Selleckchem). The following day (day 1), differentiation media 1 was changed. On day 2, differentiation media 1 was changed to differentiation media 2; MNb supplemented with 0.1 µM retinoic acid (RA) (AA4454077, Fisher scientific) and 0.5 µM smoothened agonist (SAG) (SML1314-1MG, Merck). Media 2 was changed on day 4. On day 7, differentiation media 2 was changed to differentiation media 3; MNb supplemented with 0.1 µM RA, 0.5 µM SAG, BDNF (10 ng/ml, 450-02-10UG, Thermo Fisher), and GDNF (10 ng/ml, 450-02-10UG, Thermo Fisher). On day 9, differentiation media 3 was changed to media 4; MNb supplemented with 0.1 µM RA, 0.5 µM SAG, 10 ng/ml BDNF, 10 ng/ml GDNF, and 20 µM DAPT (565770, Merck). Motor neuron progenitor spheroids were dissociated into single cells for plating on day 10 by using Accumax (00-4666-56, Thermo Fisher). Dissociated motor neuron progenitors were then plated on poly-D-lysine 50 μg/ml (A-003-E, Merck Millipore) and laminin 10 μg/ml (L2020, Merck) coated coverslips in 24-well plates or coated 6-well plates in differentiation media 4 additionally supplemented with 10 μM ROCKi. Half of media 4 was changed on day 11 (no ROCKi added). At day 14, half of media 4 was changed to media 5; MNb supplemented with 10 ng/ml BDNF, 10 ng/ml GDNF, and 20 µM DAPT. At day 16 half of media 5 was changed to media 6; MNb supplemented with 10 ng/ml BDNF, 10 ng/ml GDNF, 10 ng/ml CNTF (450-13-5UG, Thermo Fisher), and 20 µM DAPT. From day 18 onward, the cells were switched to motor neuron maturation medium; MNb supplemented with BDNF, GDNF, and CNTF (each 10 ng/ml). Media were changed every other day during the first week of maturation, days 18 to 25, and twice a week afterward by replacing half of the medium. The mature MNs were processed on day 32 of differentiation.

### Human RA-gastruloid differentiation

For RA-gastruloid induction from the H9 cell line, an adapted protocol from PMID: 39164488 was employed. In summary, 6 × 104 H9 hES cells were seeded on 0.5 μg cm−2 Vitronectin-coated 3.5 cm plate in 2 mL of Essential 8 medium with 10 µM Y-27632. On day 1, the E8 the medium was exchanged with fresh medium without any inhibitor. On day 2, the medium was replaced with Essential 8 containing 5 µM CHIR. On day 3, the medium was replaced with Essential 8 containing 500 nM RA, 5 µM CHIR and 10 µM SB431542. On day 4, cells were rinsed by 0.5 mM EDTA twice, and colonies were dissociated and broken into single cells in the Ndiff media (50 mL of media: 24.5 mL of DMEM/F12+Glutamax, 24.5 mL of Neurobasal medium, 0.25 mL of N2, 0.5 mL of B27, 0.25 mL of Glutamax) Then, 5 000 cells were seeded into Nunclon Sphera 96-Well U Bottom (Thermo Fisher, 174929) with 50 µL of NDiff 227 media containing 0.5 µM CHIR and 5 µM Y-27632. On day 5, 150 µl of NDiff media was added to each well. On day 6, 150 µl of the medium was removed and fresh 150 µl NDiff medium containing 5% Matrigel and 100 nM RA was added and maintained in a 37 °C, 5% CO2 incubator until day 9.

### Ectodermal organoid culture

Ectodermal organoids were created following the protocol published by Taroc et al. (33). Briefly, wildtype and LCCS1-KI hESCs were cultured as described above, except for using mTeSR1 medium (85850, STEMCELL technologies). After reaching 80-90% confluency, the cells were dissociated with 1 mg/ml Collagenase IV (17101046, STEMCELL Technologies) at +37℃ for 40 min. Once the cell colonies showed signs of detachment: loose edges and formation of small holes in the colonies, the Collagenase IV solution was removed, and cell colonies were carefully scraped into 5 ml of DMEM-F12 with GlutaMAX (11320033, Thermo Fisher) and washed once with DMEM/F12. The colonies were then carefully transferred to ectodermal organoid medium (1:1 DMEM/F12 with GlutaMAX and Neurobasal medium (21103049, Gibco), 40 ng/ml Fibroblast Growth Factor 2 (100-18B, Peprotech), 40 ng/ml Human Epidermal Growth Factor (E9644-0.2mg, Sigma-Aldrich), 10 ug/ml Insulin (16634-50mg, Sigma-Aldric), 0.5X B27 supplement, 0.5X N2 supplement, 1 mM ROCKi, penicillin/streptomycin), colonies were triturated 1-2 times to obtain colonies of approximately 30-50 cells and transferred to T25 flasks to culture as floating aggregates +37℃, 5% CO2 in a humidified incubator. After 24 hours, the debris was removed by change of medium. On the 3rd day, individual organoids were transferred to glass culture chamber slides (178599, Thermo Scientific). By day 6-7 the organoids started to spontaneously adhere to the glass and cells started to form migrating mats of neural crest cells. Samples were collected for immunostaining on day 9.

### RNA isolation

RA-gastruloids were collected on day 9 into TRIzol (15596018; Thermo Fisher), three gastruloids per sample, three samples were processed per genotype. Gastruloids were homogenized using the 25G needle and RNA was isolated using the standard phenol–chloroform extraction. RNA was similarly isolated from control and LCCS1 fetal fibroblasts. Fibroblasts were seeded on 6-well plates (300 000 cells) and grown overnight. Next day, both Control and LCCS1 fibroblasts were either straight collected into TRIzol reagent (15596026, Thermo Fisher, Carlsbad, California, USA) or cultured with D-actinomycin (10 µg/mL, SBR00013-1ML, Merck, Poznan, Poland) in 3 mL of growth media for 1, 3, and 9 hours. Library preparation (poly A enrichment), RNA-sequencing (NovaSeq X Plus Series: PE150,12.00 G raw data per sample), and mapping (HISAT2) (60) were performed by Novogene (Munich, Germany).

### Long range RNA sequencing

RNA isolated from the WT, control (broken PAM sequence) and LCCS1-hESCs was used for PacBio SMRT sequencing to analyze the presence of any undesired modifications or changes in length of the *GLE1* transcript. Sample purity was checked with NanoDrop (ND2000USCAN, Thermo Fisher) and the RNA integrity with Agilent 2100 Bioanalyzer. The library has been prepared from 150 µg of RNA using the SMRTbell prep kit guideline. The quality control, library preparation, and sequencing have been performed by Novogene (UK). Following the sequencing, the demultiplexed files were further processed using PacBio’s IsoSeq bioconda package (v. 4.0.0) following tool documentation provided by Pacific Biosciences. In short, any remaining primer sequences were removed from sample spesific HiFI-reads using lima tool in isoseq mode. Next, ’isoseq refine’ was used to trim Poly-A tails and to identify and remove concatamers. Isoform-level clustering was done using ’isoseq cluster2’ function, which was followed by alignment to reference genome using pbmm2, a minimap2 SMRT wrapper for PacBio data. Reference genome and annotations used were from GENCODE release M33. ’Isoseq collapse’ was used to collapse redundant transcripts into unique isoforms using recommended settings for bulk tissue IsoSeq. Pigeon PacBio Transcript Toolkit was then used to prepare the input files and then to classify transcripts against given annotations.

### RNA sequencing

Messenger RNA was purified from total RNA using poly-T oligo-attached magnetic beads. After fragmentation, the first strand cDNA was synthesized using random hexamer primers followed by the second strand cDNA synthesis. The library was ready after end repair, A-tailing, adapter ligation, size selection, amplification, and purification. The sequencing was performed by Novogene (Germany) on Illumina NovaSeq X Plus Series (PE150).

### mRNA degradation rate calculation

The methodology for calculating mRNA degradation rates was adapted from the previously described large-scale normalization approach (61). Following this method, raw sequencing reads were pre-processed to remove adapter sequences and low-quality bases. The cleaned reads were then aligned by using a standard RNA-seq aligner to generate BAM files. The BAM files were indexed to enable efficient access. Gene-level read counts were quantified using featureCounts from the Rsubread package. Reads were mapped to exonic regions and summarized at the gene level. The raw count data was then filtered to include only protein-coding genes (mRNAs) based on the gene biotype information in the GTF annotation.

To account for sequencing depth, gene length, and group-specific variations, a two-step normalization procedure was applied following the method of Viegas et al. (2023). First, standard Transcripts Per Million (TPM) normalization was performed. The TPM value for each gene was calculated using the following formula:

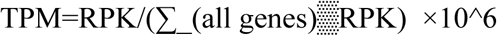

where RPK (Reads Per Kilobase) is the raw read count divided by the gene length in kilobases. Second, a set of stable normalization genes was identified to be correct for any remaining non-biological variation. For each gene, a linear regression model was fitted to its log-transformed TPM values over the four time points. A gene was classified as stable if its expression level exhibited a very low degradation rate (model slope <0.02) and the decay was highly linear (R-squared value >0.7). The final set of normalization genes consisted of the intersection of genes that met these criteria in both the Control and LCCS1 groups. The average log-transformed TPM of these stable genes was then calculated for each time point. This average value was used as a normalization factor to correct the expression data of all genes (not just the stable ones) in a time-point-specific manner. The final data was expressed as log-normalized TPM values.

For each gene that passed an initial abundance filter (at least one sample with a normalized TPM of 5 or higher), a linear regression model was fitted to its log-normalized TPM values across the time course for both the Control and LCCS1 groups separately. The beta degradation rate (β) for each gene was defined as the negative of the model’s slope. The goodness-of-fit for each model was assessed using the coefficient of determination (R-squared). Genes with an R-squared value of less than 0.5 in either group were excluded from the final analysis to ensure a high quality of fit. The mRNA half-life (t1/2) for each gene was then calculated using the formula:

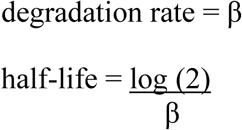

The statistical significance of the difference in mRNA degradation rates between the Control and LCCS1 groups was assessed by fitting a single linear model with an interaction term (Normalized_log_TPM ∼ Timepoint_num * Group). The p-value associated with the interaction term, documented in the P_value_Group_Comparison column, was used to determine if the slopes (and thus the beta degradation rates) were significantly different between the two groups.

### Software and Packages

All analyses were performed in R v4.4.1 (R Foundation for Statistical Computing, Vienna, Austria) and RStudio version 2024.12.1 Build 563 (Posit; Boston, MA, USA). The computational analyses were conducted using the following Bioconductor and CRAN packages for data processing and visualization: dplyr (v1.1.0), ggplot2 (v3.4.2), tidyr (v1.3.0), ggrepel (v0.9.3), Rsubread, Rsamtools (v2.16.0), GenomicFeatures (v1.52.1), rtracklayer (v1.60.1), Biostrings (v2.68.1), stringr (v1.5.0), txdbmaker (v1.0.0), GenomeInfoDb (v1.36.1), AnnotationDbi (v1.62.1), and GenomicRanges (v1.52.0). Additional packages used for downstream analysis included DESeq2, clusterProfiler, ReactomePA, org.Hs.eg.db, and enrichplot. All scripts included internal error handling, reproducibility safeguards (fixed random seeds), and local caching of annotation resources to ensure stable performance without reliance on persistent internet access. All packages were run with their default parameters unless otherwise specified.

### Polysome profiling

hESC-derived spinal motor neurons were differentiated in 6-well plates (two independent differentiations) and harvested for polysome profiling. Prior to harvesting, cells were washed with ice-cold PBS supplemented with cycloheximide (100 µg/mL) to stabilize translating ribosomes. Cells were lysed on ice in polysome lysis buffer containing 20 mM Tris-HCl (pH 7.5), 150 mM NaCl, 10 mM MgCl2, 1% NP-40, 1% sodium deoxycholate, 1 mM DTT, DNase I, cycloheximide, and protease inhibitors. Lysates were homogenized by repeated passage through a 26-gauge needle and clarified by centrifugation at 10,000 × g for 10 min at 4°C.

Clarified lysates were subjected to sucrose gradient ultracentrifugation for polysome separation. Fractions corresponding to polysome-associated RNA were collected and pooled. RNA was isolated from pooled polysomal fractions using TRIzol reagent according to the manufacturer’s instructions. Following RNA purification, cDNA was synthesized and transcript abundance of genes of interest was quantified by RT-qPCR.

### Reverse transcription-quantitative polymerase chain reaction (RT-qPCR)

The cDNA was synthesized using the SuperScript IV Reverse Transcriptase (18090050, Thermo Fisher) and diluted in nuclease free water. For the qPCR reaction, 2.5 µl of cDNA were mixed with 1 µl each of forward and reverse primer (Supplementary table 2) along with 1.5 µl of PCR water and 5 µl SensiFAST™ SYBR® No-ROX Kit (BIO-98020, Bioline. qPCR was carried out at CFX384 Touch Real-Time PCR Detection System (Bio-Rad). qPCR was performed with at least three biological replicates in two technical replicates. The gene expression was normalized to 20 pg of Luciferase mRNA spike-in control (62) (L4561, Promega) since the constitutive gene expression was questionable, and the relative gene expression was calculated using the 2^−ΔΔCT^ method (63).

### Immunostaining

Cells were seeded or differentiated on coated glass coverslips – patient amniotic cells: fibronectin (1030-FN, R&D Systems), hESCs: matrigel, postmitotic and mature MNs: poly-D-lysine 50 μg/ml (A-003-E, Merck Millipore) and laminin 10 μg/ml (L2020, Merck).

### hESCs and patient amniotic cell samples

Cells were fixed with 4% PFA in PBS for 20 min at room temperature and washed twice with wash buffer (0.2% BSA in PBS + 0.1% Tween 20). The cells were permeabilized with 0.1% Triton X-100 + 50 mM glycine for 15 minutes, washed with wash buffer three times while on horizontal shaker, blocked with 5% FBS in 0.1% Triton X-100 in PBS to block unspecific binding and incubated with primary antibodies (Supplementary table 3) diluted in the blocking buffer overnight at 4°C. Cells were washed with the wash buffer three times again before incubation with the secondary antibodies diluted in the blocking buffer for 90 min. Following another washing step, the nuclei were stained with Hoechst for 8 min. After washing the cells with PBS, the cover slips were mounted on glass slides using the Epredia™ Immu-Mount™ (10622689, Fisher Scientific)

### Postmitotic and mature MNs

Neurons were gently rinsed in PBS once before fixation in 4% PFA for 10 min. After fixation, cells were washed by PBS 3x and penetrated by 0.2% Triton X-100. Cells were blocked in 5% BSA in PBST. Primary antibodies were diluted in the blocking buffer and incubated with the cells overnight at +4 °C. Next day, cells were washed in PBS trice, before incubating them with secondary antibodies for 90 min. Cells were washed in PBS trice and the nuclei were counterstained with Hoechst for 8 min. After washing the cells with PBS, the coverslips were mounted on glass slides using the Epredia™ Immu-Mount™.

### Fibroblasts

Cells were seeded and grown on fibronectin-coated coverslips until desired confluence. Cells were fixed with 4% PFA in PBS for 20 min at room temperature and washed twice with wash buffer (0.2% BSA in PBS + 0.1% Tween 20). The cells were permeabilized with 0.1% Triton X-100 + 50 mm Glycine for 15 min, washed with wash buffer three times while on a horizontal shaker, blocked with 5% FBS in 0.1% Triton X-100 in PBS to block unspecific binding, and incubated with primary antibodies) diluted in the blocking buffer overnight at 4 °C. Cells were washed with the wash buffer three times again before incubation with the secondary antibodies diluted in the blocking buffer for 90 min. Following another washing step, the nuclei were stained with Hoechst for 10 min. After washing the cells with PBS, they were mounted on glass slides using the Epredia™ Immu-Mount™ (9990412; Thermo Fisher, Landsmeer, Netherlands). Optionally, phalloidin staining has been added after the secondary antibody staining by diluting the phalloidin in PBS and staining for 2 h.

### Western blot

Cells were washed with ice cold PBS twice and lysed using 1x RIPA cell lysis solution supplemented with Pierce™ Protease Inhibitor Tablets, EDTA-free (A32965, Thermo Fisher) and Pierce™ Phosphatase Inhibitor Mini Tablets (A32957, Thermo Fisher). The homogenate was pushed through the 25G needle several times, centrifuged at 10 000 RPM for 15 min, and the supernatant was collected for analysis. Protein quantity estimation was performed using Pierce™ BCA Protein Assay Kit (23225, Thermo Fisher). In summary, 3.5 µg of mature MN cell lysate was used for analysis. Protein was loaded on to in-house prepared SDS-PAGE separating gels and separation was performed at 110V at room temperature. Transfer was carried out using the wet blot transfer onto the nitrocellulose membrane. The membrane was blocked with 5% BSA solution for 2 hours and all the primary antibodies (Supplementary table 4) were incubated overnight at +4 °C, while secondary antibodies were incubated for 90 min at room temperature. The detection was performed using the WesternBright ECL HRP substrate on Bio-Rad ChemiDoc™ Imaging System. The proteins were either normalized to the total protein load quantified by the Ponceau S (78376, Merck) staining, or to β-actin after the antibodies were stripped away by the Restore™ PLUS Western Blot Stripping Buffer (46430, Thermo Fisher) for 10 min at room temperature and re-blocking and re-staining.

### Cellular thermal shift assay

The thermal stability (64) of GLE1 protein was tested in WT and LCCS1-hESCs. The hESCs were cultured on 6 cm diameter plates coated with Matrigel, three plates per genotype. Plates were washed with ice-cold PBS twice before scraping the adherent cells off the dishes into PBS using a cell scraper. The cell suspension was equally aliquoted into PCR tubes, with 100 μl of suspension per tube. The cell suspensions were heated individually for 5 min in the C1000 Touch Thermal Cycler (Bio Rad) and allowed to cool down in 10 °C for 5 min. The heated cell suspensions were lysed by three cycles of freeze-thawing with liquid nitrogen and a 37 °C water bath. The cell lysate mixture was centrifuged at 15 000 RPM for 10 min at +4 °C. About 80 μl of supernatant (non-aggregated soluble proteins) per sample were removed, mixed with loading buffer, and the mix heated at 95 °C–100 °C for 10 min before WB analysis.

### Oligo dT hybridization

Fluorescent *in situ* hybridization of the poly(A)+ RNA was performed on the cells on coverslips to analyze the mRNA abundance and distribution. Cells were fixed with 4% PFA in PBS for 20 min at room temperature, washed in PBS twice, permeabilized with ice cold methanol for 10 min and then incubated in 70% ethanol at +4 °C overnight. On the next day, the cells were incubated for 5 min at room temperature in wash buffer (10% formamide in 2× saline sodium citrate (SSC)). Then, the cell were treated by 200 μl of hybridization buffer (10% formamide (AM9342, Thermo Fisher), 2× SSC, 1 mg/ml yeast tRNA (AM7119, Thermo Fisher) and 10% Dextran sulfate (J14489, Thermo Fisher) containing 100 nM probe (5’-Cy3-oligo-dT(30), IDT) , in a humidified chamber sealed with parafilm and incubated in the dark at 37°C overnight. After hybridization, the coverslips were washed twice in wash buffer in the dark at 37°C, for 30 min each wash, and then washed briefly in 1× PBS. Hoechst (10 min) was used as a DNA counterstain.

### RNA synthesis

Global RNA synthesis was analyzed in mature MNs through metabolic labeling. Cells were treated with 1 mM ethynyl uridine (EU) for 1 h before fixation with 4% PFA in PBS for 20 min. Cells were washed and permeabilized with 0.5% Triton X-100 in PBS and click chemistry was performed according to the Click-iT™ RNA Alexa Fluor™ 594 Imaging Kit (C10330, Thermo Fisher) instructions.

### Protein synthesis

Global protein synthesis was analyzed through metabolic labeling. Cells were treated with 20 µM O-propargyl-puromycin for 30 min before fixation with 4% PFA in PBS for 20 min. Cells were washed and permeabilized with 0.5% Triton X-100 in PBS and click chemistry was performed according to the Click-iT™ Plus OPP Alexa Fluor™ 594 Protein Synthesis Assay Kit (C10457, Thermo Fisher) instructions.

### Proliferation assay

EdU labelling was performed using Click-iT EdU Alexa Fluor 488 Imaging Kit (C10337, Thermo Fisher) according to the manufacturer’s instructions. Cells were incubated with 10 µM EdU in the culture medium in a 37°C incubator (5% CO_2_) for 2 h before fixation with 4% PFA in PBS for 30 min.

### O-linked protein glycosylation

Fibroblasts were seeded (40 000 cells) and grown on fibronectin-coated coverslips overnight. Cells were treated with 50 µM GalNAz (C33365, Thermo Fisher, Bleiswijk, Netherland) or 50 µM GlcNAz (C33367, Thermo Fisher, Bleiswijk, Netherland) for 72 hours. Cells were washed 2x with PBS, fixed with 4% PFA in PBS for 20 min, and permeabilized in 0.1% Triton X-100 in PBS for 5 min. Subsequently, click chemistry was performed with 250 µL of working solution: 10 µM AF488-alkyne (A10267, Thermo Fisher, Carlsbad, California, USA), 1 mM CuSO4, 200 µM THPTA ligand, and 5 mM sodium ascorbate. Cells were counterstained with 1:1000 Hoechst diluted in PBS for 10 min, washed in PBS, and mounted with the Epredia™ Immu-Mount™.

### Image acquisition

For intensity measurements, the cells were imaged with the Zeiss Axio Imager equipped with Hamamatsu Orca Flash 4.0 LT B&W camera and using the EC Plan Neofluar 40x/1.3 and 63x/1.4 objectives. To visualize the stress granules and NF-L inclusions, cells were imaged by the Andor Dragonfly spinning disc microscope (z-stack: 0.25 µm per step) with Zyla 4.2 sCMOS camera and Plan Apo VC 60x/1.2 objective.

For ectodermal organoid samples, images of the entire adherent organoids were obtained using Leica Leica M 205 FCA stereomicroscope equipped with Leica K5 B&W sCMOS camera 1x PlanApo stereo objective. High-resolution images from different positions of the adhered organoids were obtained with Zeiss LSM880 confocal microscope equipped with 40x/1.40 Plan-Apochromat oil objective.

### Image analysis

Cells have been segmented by CellProfiler 4.2.1 either to measure the mean fluorescence intensity of target structures in subcellular compartments or to quantify the number of stress granules (speckle-type objects per cell) or the number of NF-L inclusions.

For organoid samples, ImageJ (1.54p) and plugin StarDist (65) was used. After segmentation of all nuclei, the condensed nuclei of dead cells were removed by size exclusion. Mean intensity values in the Sox2 and Sox9 channel were used to calculate the ratio of Sox2 to Sox9 signal in 100 – 801 nuclei per sample. The percentage of Sox2 and Sox9 positive nuclei was determined from 3 samples of each genotype by thresholding the mean intensity value.

### Statistics and reproducibility

Statistical analyses were performed using GraphPad Prism software (GraphPad, version 10) or R (2022.12.0 Build 353). Statistical significance was determined by the specific tests indicated in the corresponding figure legends. Only two tailed tests were used. No statistical methods were used to predetermine sample sizes. Data collection and analysis were not performed blinded.

## Data availability

The RNA sequencing results (PRJNA1183003, PRJNA1467873, and PRJNA1468682) have been deposited in NCBI’s Sequence Read Archive (SRA).

## Ethics statement

The present study protocol was evaluated, reported, and approved by the Institutional Review Board of Helsinki University Hospital (approval No. HUS/6254/2023) in accordance with local policies and the ISSCR Guidelines for Stem Cell Research and Clinical Translation. Because the experimental approach did not involve the fertilization of gametes or the intention to generate a human embryo, it conforms to Category 1B protocols. Studies involving previously collected human fetal material follow the Declaration of Helsinki (WMA, 2013) and informed consent procedures, and are in line with Directive 2004/23/EC.

## Disclosure

All authors have no potential conflicts of interest.

## Funding

Research Council of Finland. Grant Number: 348906; Jane ja Aatos Erkon Säätiö; Helsinki Institute of Life Science, Helsingin Yliopisto.

## Author contributions

Conceptualization: Zarybnicky T, Pakkasjärvi N, Kuure S; Data curation: Zarybnicky T, Lindfors S, Väänänen J; Formal analysis: Zarybnicky T, Mäkinen OJM, Balatsou A, Yalcin S, Lindfors S, Metso S, Gregorová P, Väänänen J, Rekonen L, Bůtorová K; Funding acquisition: Zarybnicky T, Kuure S; Investigation: Zarybnicky T, Lindfors S, Balatsou A; Methodology: Zarybnicky T, Lindfors S, Sartori Maldonado R, Gregorová P, Sarin PL, Tyynismaa H, Wartiovaara K; Project administration: Kuure S; Resources: Wartiovaara K, Tyynismaa H, Kuure S; Writing -original draft: Zarybnicky T, Kuure S; Writing -review & editing: Zarybnicky T, Balatsou A, Sartori Maldonado R, Tyynismaa H, Wartiovaara K, Pakkasjärvi N; Study supervision: Kuure S.

## Acknowledgements

We thank the families who have granted human material for biomedical studies, Helsinki Biobank for distributing the tissues, and HUSLAB Genetics laboratory. The following core facilities of Biocenter Finland supported by the Helsinki Institute of Life Science are acknowledged: the GM-Unit Core Facility for providing mouse embryonic fibroblasts, the Biomedicum Imaging Unit for microscopy services, and the University of Helsinki Flow Cytometry Unit for technical assistance. This work was supported by Academy of Finland (348906), Jane and Aatos Erkko Foundation, and University of Helsinki funds.

## Notes

https://www.ncbi.nlm.nih.gov/bioproject/1467873

### Competing Interest Statement

The authors have declared no competing interest.

