## Supplemental info for "Human models of GLE1-associated LCCS1 reveal neural crest deficiency and multisystem developmental failure accompanied by altered RNA metabolism"

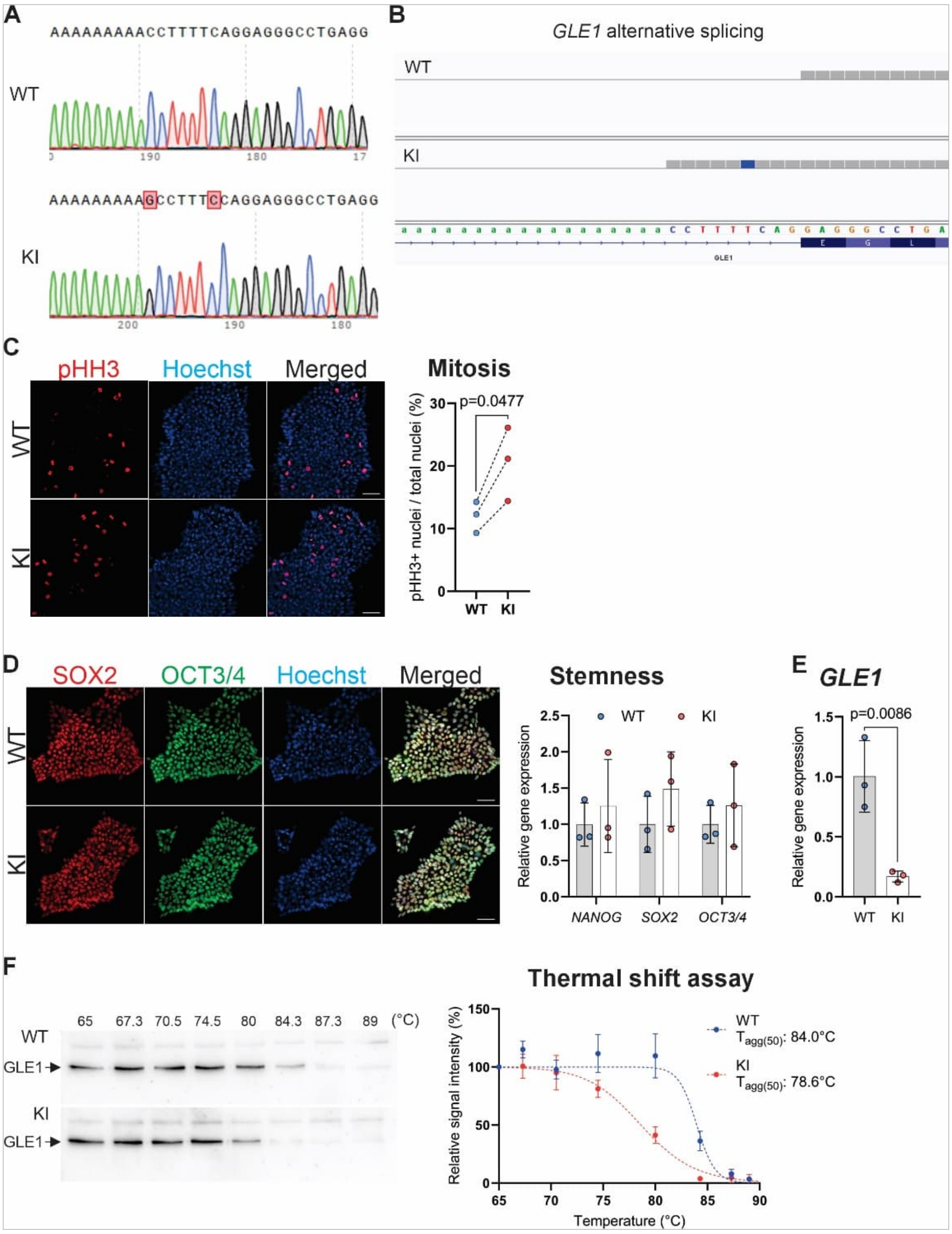


**Supplementary Figure 1. Characterization of LCCS1 hESC model**

A) Sanger sequencing chromatographs of the targeted genomic sequence in wild type (WT, top) and knock-in homozygote (KI, bottom) hESCs. The result demonstrates the successful CRISPR CAS12-mediated insertion of the A-to-G nucleotide at the intron/exon boundary of human *GLE1* gene.

B) Long-read sequencing of the wild type (WT) and c.432-10A>G variant carrying knockin cells (KI). The desired nine nucleotide insertion is only detected in the KI hESCs.

C) Representative images shown for immunofluorescent labeling of phospo-Histone H3 (pHH)+ cells depicting all wild type (WT) and knockin (KI) human embryonic stem cells (hESCs) in mitosis. Quantification of proliferative cells positive for pHH3 (red) among all nuclei visualized by Hoechst (blue). The percentage of pHH3-positive cells in each independent experiment (over 5000 cells per experiment, three independent experiments) is presented. Scale bar: 50 µm. Paired t-test.

D) Immunostaining of classical stemness regulators SOX2 (red) and OCT3/4 (green) in wild type (WT) and knock-in homozygote (KI) hESCs. Scale bar: 50 µm.

B) Gene expression analysis of stemness regulators (*NANOG*, *SOX2*, *OCT3/4*) by reverse transcriptase-quantitative PCR (RT-qPCR) in three different yet comparative passages of wild type (WT) and knock-in (KI) hESC.

E) Gene expression analysis of *GLE1* in wild type (WT) and knockin (KI) hESC cells by RT-qPCR. Three independent yet comparative passages, student's t test.

F) Analysis of GLE1 protein stability in wild type (WT) and knockin (KI) hESCs using the cellular thermal shift assay. Three independent experiments were carried out, and non-linear regression was plotted to calculate the aggregation temperature ($T_{agg(50)}$) for GLE1 in both genotypes. Lower aggregation temperature indicates decreased thermal stability of GLE1, which may reflect altered protein folding, increased aggregation propensity, and/or reduced functional integrity.

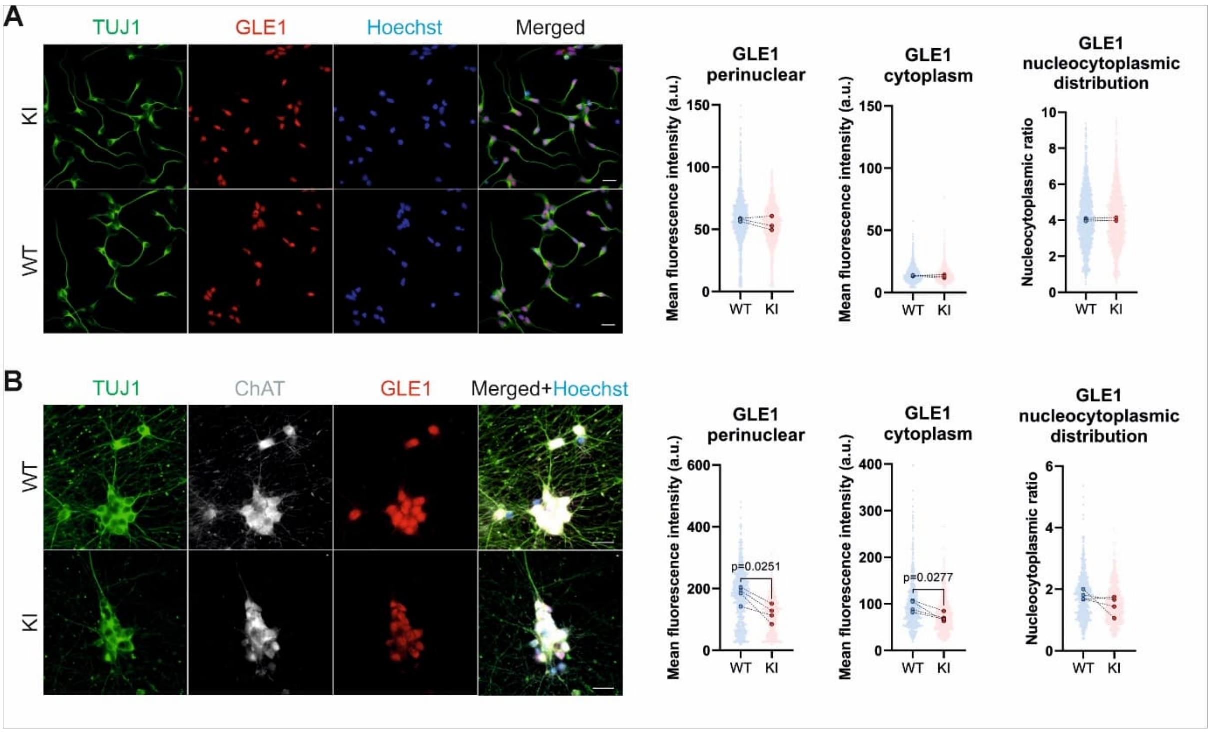


Supplementary Figure 2. **GLE1 distribution in differentiating motor neurons**

A) Immunostaining, quantification, and subcellular distribution of GLE1 (red) in postmitotic (D11) wild type (WT) and LCCS1 knockin (KI) motor neurons (MNs), co-stained with TUJ1 (green) visualizing the entire neuron structure. Individual data points generated from each cell are plotted (500 cells per differentiation; three independent differentiations), as well as median values from each experiment (large dots) are shown. Scale bar: 20 µm. Paired t-test.

B) Immunostaining, quantification, and subcellular distribution of GLE1 (red) in mature (D32) wild type (WT) and LCCS1 knockin (KI) MNs, co-stained with neuronal marker TUJ1 (green) and motor neuron marker ChAT (white). Individual data points generated from each cell are plotted (150 cells per differentiation, four independent differentiations), as well as median values from each experiment (large dots) are shown. Scale bar: 20 µm. Paired t-test.

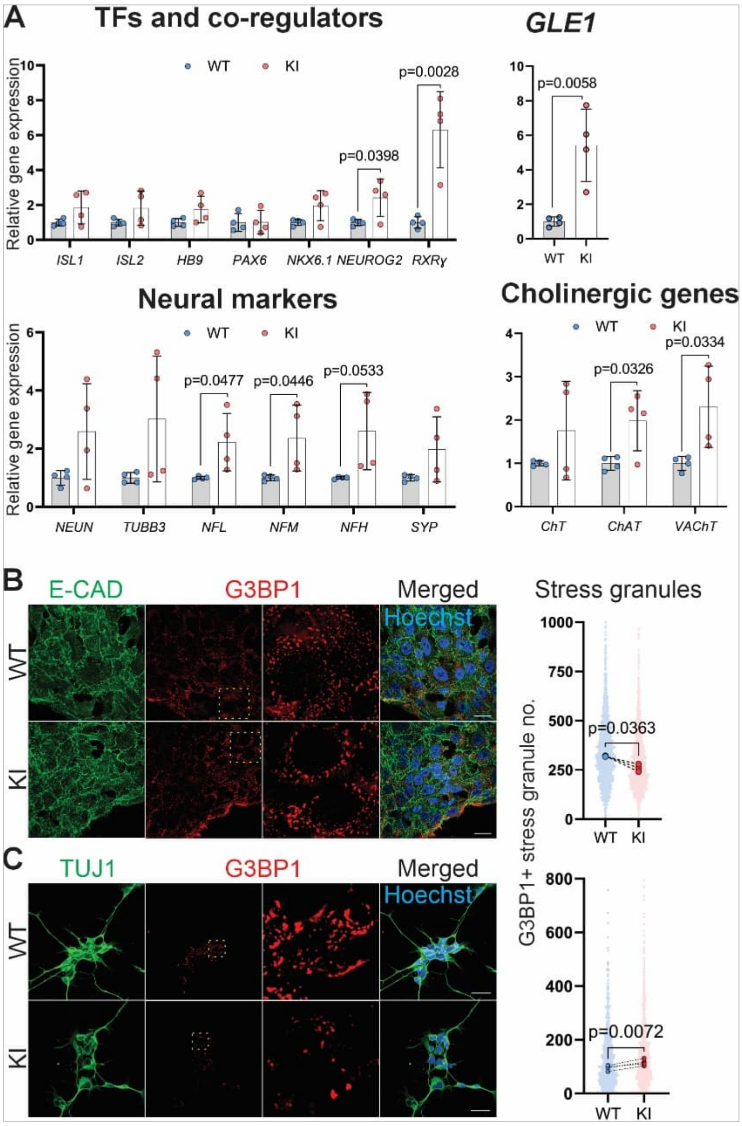


Supplementary **fi**gure 3. Analysis of motor neuron master regulators **and stress granule formation**

A) RT-qPCR quantification of transcription factors (TFs) and their co-regulators, neural markers, cholinergic genes and *GLE1* in WT and KI mature motor neurons (MNs). Individual data points from four independent differentiations are plotted, as well as mean ± SD. Student's t test.

G3BP1-positive stress granule counts after hyperosmotic shock during MN differentiation. Hyperosmotic shock (400 mM sorbitol, 60 min) induced to B) hESCs and C) postmitotic MNs followed by quantification of immunostained G3BP1-positive stress granules (red, zoomed in areas in yellow squares) co-stained with E-cadherin (green) in hESCS and with TUJ1 (green) in postmitotic MNs. Individual data points generated from each cell are plotted (500 (hESCs) and 400 (MNs) cells per experiment, three independent experiments), as well as median values from each experiment (large dots). Scale bar: 20 µm. Paired t-test.

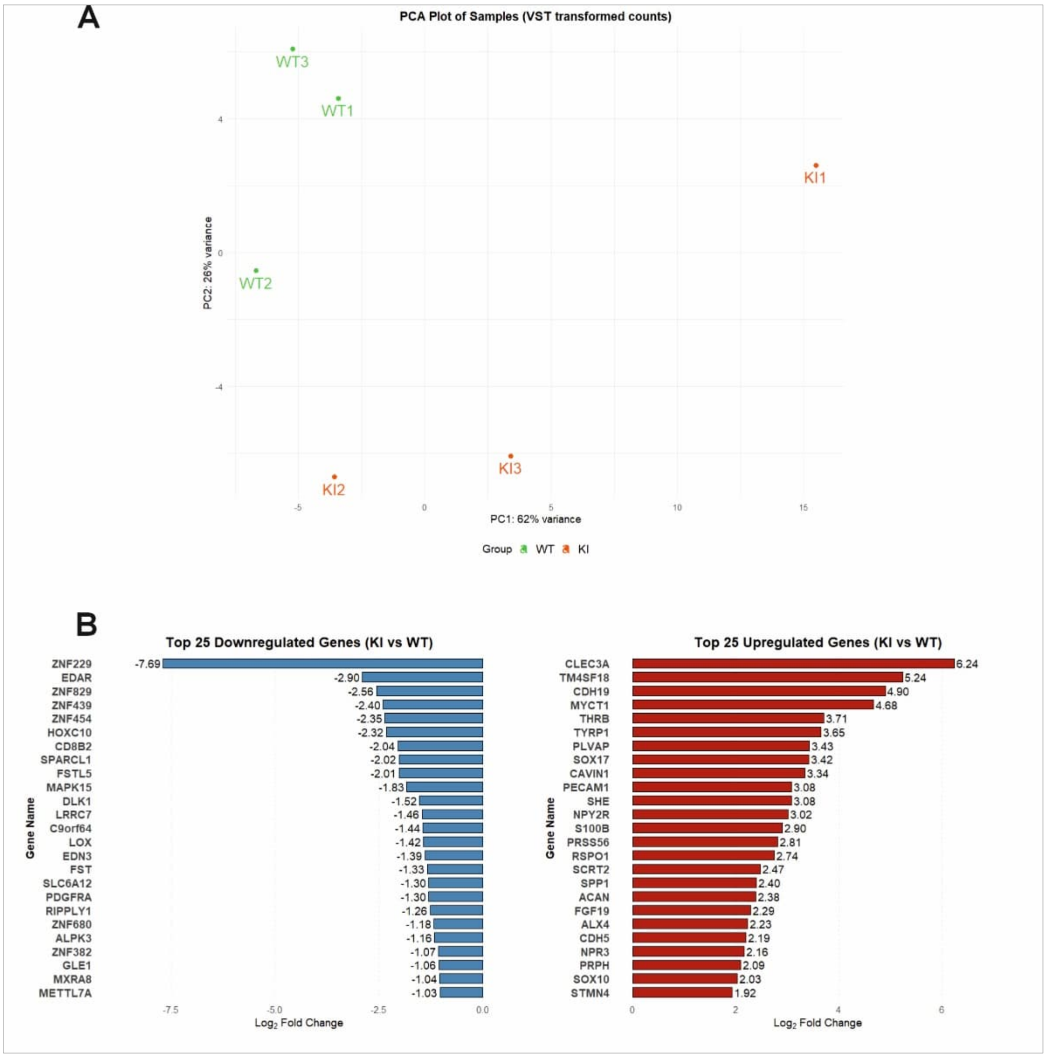


Supplementary figure 4. **Transcriptional characterization of LCCS1** gastruloids.

A) Principal component (PC) analysis of the transcriptomic analysis of gene expression differences in gastruloids derived from wild type (WT, n=3) and LCCS1 knockin (KI, n=3) human embryonic stem cells. Points represent the transcriptome of WT gastruloids in green and LCCS1 in orange. PC1 represents principal component one, and PC2 represents principal component two, both of which explain the first and the second higher variance among genotypes.

B) Identities and $\log_2$ fold changes of top 25 down- (blue) and upregulated (red) transcripts in LCCS1 gastruloids.

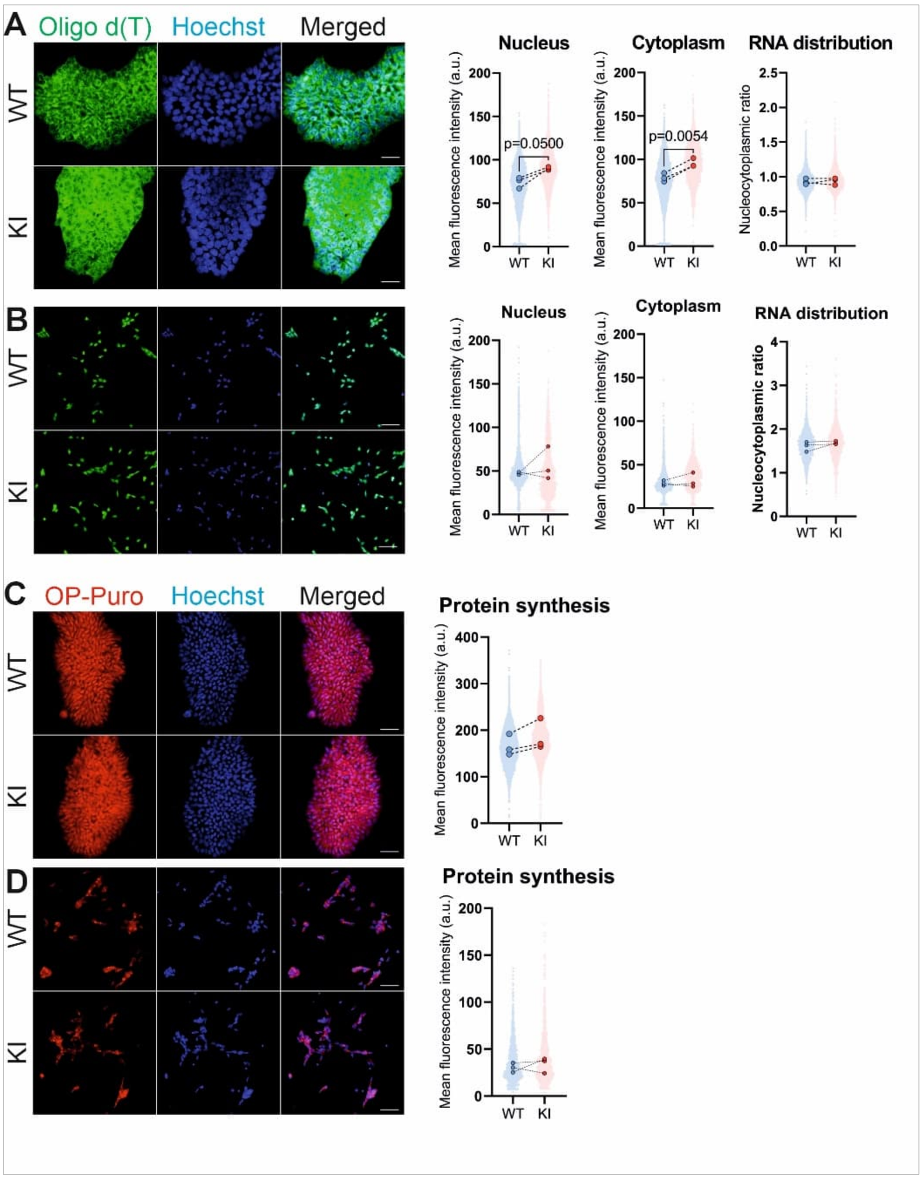


Supplementary **fi**gure 5. mRNA **distribution** and **protein synthesis in hESCs and postmitotic MNs**. Oligo d(T) probe fluorescent *in situ* hybridization (FISH) to detect the poly(A)+ RNA and quantify its subcellular abundance and nucleocytoplasmic distribution in wild type (WT) and LCCS1 knockin (KI) A) human embryonic stem cells (hESCs) and B) postmitotic motor neurons (MNs). Individual data points generated from each cell are plotted (2000 cells per experiment, three independent experiments), as well as median values from each experiment (large dots). Scale bar: 50 µm. Paired t-test.

Quantification of protein synthesis during the MN differentiation via O-propargyl-puromycin metabolic labelling (20 µM OP-Puro, 30 min pulse) in C) hESCs and D) postmitotic MNs. Individual data points generated from each cell are plotted (4000 cells per experiment, three independent experiments), as well as median values from each experiment (large dots) shown. Scale bar: 50 µm.

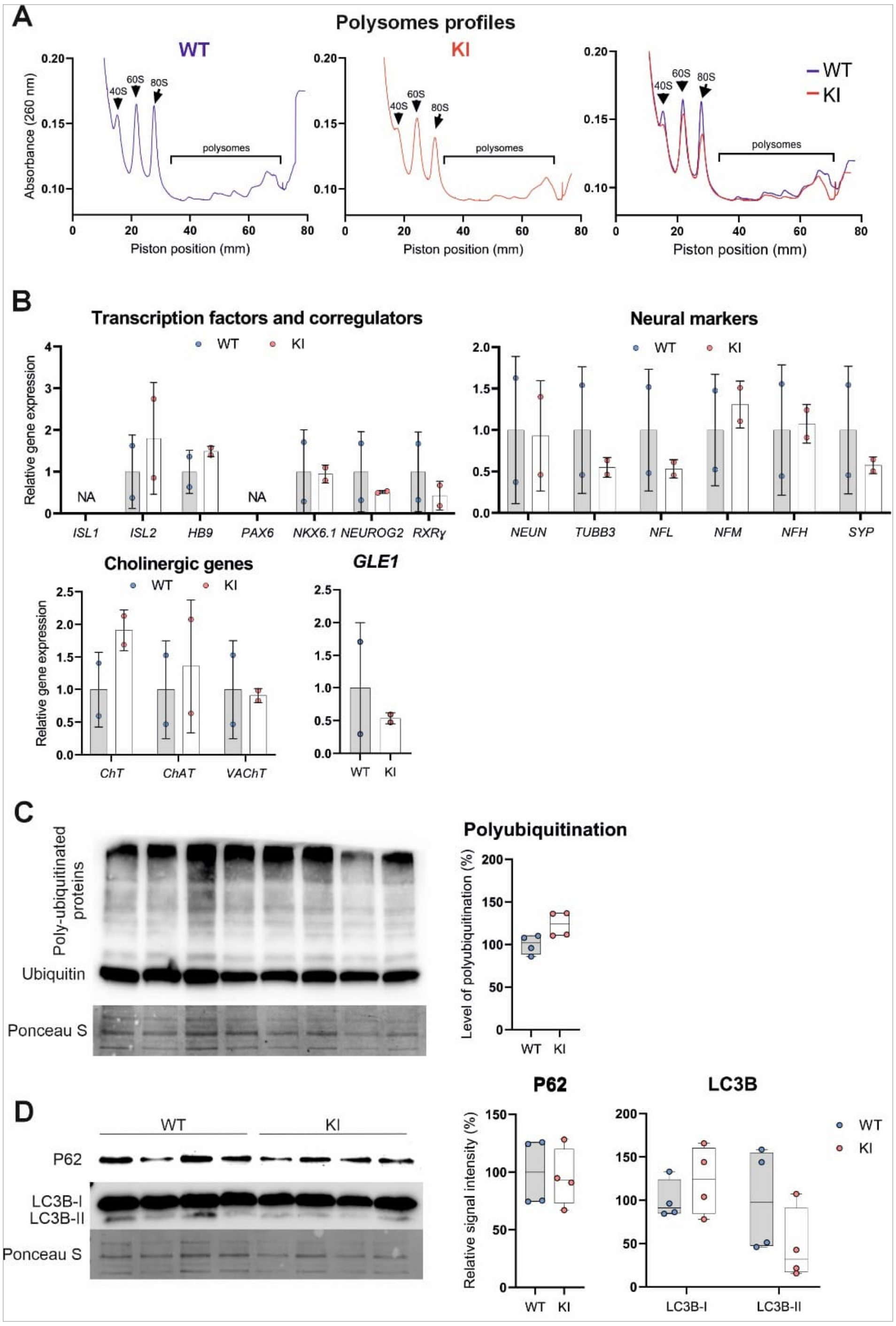


Supplementary figure 6. Protein translation and turnover in mature LCCS1 motor neurons. Representative polysome profiles showing ribosomal abundance (measured by 260-nm absorbance along a sucrose gradient) in mature wild type (WT, blue), LCCS1 variant carrying knockin (KI, red) motor neurons and their overlay.

B) Quantification of major functional genes from the sucrose gradient polysomal fraction of WT and KI mature motor neurons by RT-qPCR. Individual data points from two independent differentiations are plotted and mean ± SD shown. Student's t-test.

C) Analysis of ubiquitin homeostasis in WT and KI mature motor neurons by Western blotting. Quantification of polyubiquitinated protein amount is presented as a box plot with whiskers, including individual data points from each differentiation (four independent differentiations).

D) Measurement of autophagy-related proteins P62 and LC3B-I/II abundance and their quantification in WT and KI mature motor neurons. Data are presented as a box plot with whiskers, including individual data points from each differentiation (four independent differentiations).

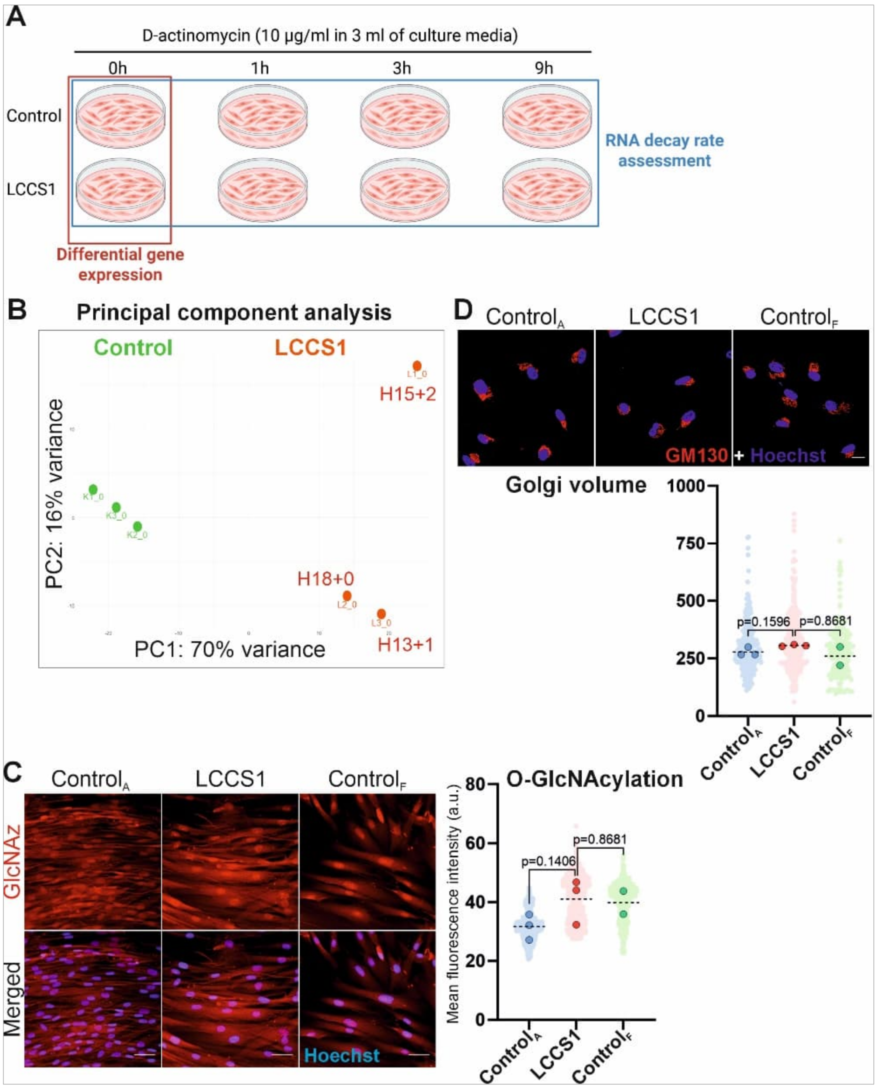


Supplementary figure 7.

A) Schematics of the RNA sequencing experiment designed to compare the transcriptome in non-treated and transcription-inhibited control and LCCS1 patient fibroblasts. For transcription inhibition, fibroblasts were treated with 10 µg/mL D-actinomycin for 1, 3, and 9 hours followed by mRNA isolation for bulk RNA sequencing.

B) Principal component (PC) analysis of the transcriptome in non-treated control (green) and LCCS1 patient (orange) cells. PC1 and PC2 explain the first and second higher variance among genotypes.

C) Metabolic labelling of patient fibroblasts with 50 µM GlcNAz (red) for 72 hours followed by the click chemistry to quantify the level of O-GlcNAcylation in control and LCCS1 derived fibroblasts. Individual data points generated from each cell are plotted (300 cells per patient, three adult control cell lines, 3 LCCS1 patient cell lines, 2 fetal control cell lines), and median values (large dots) from each patient shown. Scale bar: 50 µm.

D) Maximum projection images of control and LCCS1 patient fibroblasts immunostained with GM130 (red) as a Golgi marker. Nuclei are visualized by Hoechst (blue). Volumetric analysis of the Golgi area positive for GM130 via Imaris software. Individual data points generated from each cell are plotted (75 cells per patient, three adult control cell lines, 3 LCCS1 patient cell lines, 2 fetal control cell lines), and median values (large dots) from each patient shown. Scale bar: 20 μm.

Supplementary table 1

| Oligo | Sequence | Manufacturor |
|---|---|---|
| crRNA | AGGAGGGCCTGAGGCTATGG (PAM: TTTC) | IDT |
| ssDNA | gttgcagtgagctgcactccagcctggcgacagagagagattccatcttaaaaaaaaaaaaaaaaaaaaaaGCCTTTCCAGGAGGGCCTGAGGCTATGGCAGGAGGAGCAGGAGAGGAAGGTGCAAGCCCTCTCGGAGATGGCATCTGAACAACTG | IDT |
| *GLE1* Forward | GTAGCCCGCAGGGGTAAATG | IDT |
| *GLE1* Reverse | TCCTCTTGGGAAAGAGTTCCA | IDT |

Supplementary table 1**: Introduction of the** *GLE1*$_{FinMajor}$ variant

Oligonucleotide sequences of the crRNA and ssDNA (red G: introduced A-to-G substitution, blue C: TTTC PAM sequence broken to TTCC) repair template, used to introduce the the *GLE1*$_{FinMajor}$ mutation into the genome of hESCs, and the sequences of forward/reverse primers used for sequencing and genotyping the modified hESC colonies.

Supplementary table 2

| Oligo | Sequence | Manufacturor |
|---|---|---|
| *OCT3/4* Forward | TTGGGCTCGAGAAGGATGTG | IDT |
| *OCT3/4* Reverse | GTGAAGTGAGGGCTCCCATA | IDT |
| *SOX2* Forward | GCCCTGCAGTACAACTCCAT | IDT |
| *SOX2* Reverse | TGCCCTGCTGCGAGTAGGA | IDT |
| *NANOG* Forward | CTCAGCCTCCAGCAGATGC | IDT |
| *NANOG* Reverse | TAGATTTCATTCTCTGGTTCTGG | IDT |
| *LHX3* Forward | CGATGCTGCTGGAAACGG | IDT |
| *LHX3* Reverse | GTCGTCCTTGCAGTAAACGC | IDT |
| *HB9* Forward | GTCCACCGCGGGCATGATCC | IDT |
| *HB9* Reverse | TCTTCACCTGGGTCTCGGTGAGC | IDT |
| *ISL1* Forward | TGAAATGTGCGGAGTGTAATCAGTATTTGGAC | IDT |
| *ISL1* Reverse | CACACAGCGGAAACACTCGATGTG | IDT |
| *RXRγ* Forward | GACTGCCTCATTGACAAGCGTC | IDT |
| *RXRγ* Reverse | GACCACTGGTAGCACATTCTGC | IDT |
| *NKX6.1* Forward | GGGCTCGTTTGGCCTATTCGTT | IDT |
| *NKX6.1* Reverse | CCACTTGGTCCGGCGGTTCT | IDT |
| *ISL2* Forward | AGAGACGGGAAGACCTACTGCA | IDT |
| *ISL2* Reverse | AGCACTCGATGTGGTACACGCT | IDT |
| *PAX6* Forward | TGGGCAGGTATTACGAGACTG | IDT |
| *PAX6* Reverse | ACTCCCGCTTATACTGGGCTA | IDT |
| *NEUROG2* Forward | CAAGCTCACCAAGATCGAGACC | IDT |
| *NEUROG2* Reverse | AGCAACACTGCCTCGGAGAAGA | IDT |
| *ChAT* Forward | CATGAAGCAATACTATGGGCTCTTCTCCTC | IDT |
| *ChAT* Reverse | GACGGCGGAAATTAATGACAACATCCAAG | IDT |
| *CHT* Forward | AAAAGCCGTGGCTGGGAACTGT | IDT |

| | | |
|---|---|---|
| *CHT* Reverse | GCACTTGAGCATAGGTGGCTGA | IDT |
| *NEUN* Forward | TACGCAGCCTACAGATACGCTC | IDT |
| *NEUN* Reverse | TGGTTCCAATGCTGTAGGTCGC | IDT |
| *TUBB3* Forward | GCAACTACGTGGGCGACT | IDT |
| *TUBB3* Reverse | CGAGGCACGTACTTGTGAGA | IDT |
| *NEFL* Forward | ATGAGTTCCTTCAGCTACGAGC | IDT |
| *NEFL* Reverse | CTGGGCATCAACGATCCAGA | IDT |
| *NEFM* Forward | ACAACCACGACCTCAGCAGCTA | IDT |
| *NEFM* Reverse | GTTGAGGAGGTCCTGGTATTCG | IDT |
| *NEFH* Forward | CTGAGGAACACCAAGTGGGAGA | IDT |
| *NEFH* Reverse | TCCGACACTCTTCACCTTCCAG | IDT |
| *SYP* Forward | TCGGCTTTGTGAAGGTGCTGCA | IDT |
| *SYP* Reverse | TCACTCTCGGTCTTGTTGGCAC | IDT |
| *GLE1* Forward | CACGGGGGATCAAAGTGGAA | IDT |
| *GLE1* Reverse | CGCTTCAGTTGTTCAGATGCC | IDT |
| *Luciferase* Forward | GCGCGGAGGAGTTGTGTT | IDT |
| *Luciferase* Reverse | TCTGATTTTTCTTGCGTCGAGTT | IDT |

Supplementary table 2: List of RT-qPCR primers

Supplementary table 3

| Antibody | Host | Dilution | ID | Manufacturor |
|---|---|---|---|---|
| Sox2 | Goat | 1:200 | AF2018 | Biotechne |
| Oct3/4 | Mouse | 1:200 | SC5279 | Santa Cruz |
| GLE1 | Rabbit | 1:500 | PA5-21816 | Thermo Fisher |
| E-Cadherin | Goat | 1:500 | AF748 | R&D Systems |
| pHH3 (pSer10) | Rabbit | 1:500 | 06-570 | Millipore |
| ISL1/2 | Mouse | 1:100 | 39.4D5 | DSHB |
| OLIG2 | Rabbit | 1:500 | AB9610 | Merck |
| TUJ1 | Mouse | 1:500 | 801201 | BioLegend |
| TUJ1 | Rabbit | 1:1000 | AB18207 | Abcam |
| ChAT | Goat | 1:100 | AB144P | Merck |
| HB9 | Mouse | 1:100 | 81.5C10 | DSHB |
| ISL1 | Rabbit | 1:500 | AB20670 | Abcam |
| FoxP1 | Mouse | 1:500 | AB32010 | Abcam |
| G3BP1 | Rabbit | 1:500 | 13057-2-AP | Thermo Fisher |
| Alexa Fluor® 488 AffiniPure™ Donkey Anti-Goat IgG (H+L) | Donkey | 1:400 | 705-546-147 | Jackson ImmunoResearch |
| Donkey anti-Mouse IgG (H+L) Alexa Fluor™ 568 | Donkey | 1:400 | A10037 | Thermo Fisher |
| Donkey anti-Rabbit IgG (H+L) Alexa Fluor™ Plus 647 | Donkey | 1:400 | A32795 | Thermo Fisher |

**Supplementary table 3: List of antibodies used for** immunostaining

Supplementary table 4

| Antibody | Host | Dilution | ID | Manufacturor |
|---|---|---|---|---|
| GLE1 | Rabbit | 1:1000 | 26466-1-AP | Thermo Fisher |
| Ubiquitin | Rabbit | 1:1000 | #3933 | Cell Signalling |
| P62 | Mouse | 1:1000 | AB56416 | Abcam |
| LC3B | Rabbit | 1:1000 | NB600-1384 | Novus Biologicals |
| NF-L | Mouse | 1:1000 | SC390732 | Santa Cruz |
| NF-M | Mouse | 1:1000 | RMO270 | Thermo Fisher |
| NF-H | Mouse | 1:1000 | 801701 | Biolegend |
| Anti-rabbit-HRP | Sheep | 1:5000 | A16172 | Thermo Fisher |
| Anti-mouse-HRP | Goat | 1:5000 | **A16066** | Thermo Fisher |

Supplementary table 4: **List of antibodies used for WB**
